# Super-resolution complexome of human mitochondria elucidates translocase, morphology and OXPHOS networks

**DOI:** 10.64898/2026.05.29.728872

**Authors:** Uwe Schulte, Alexander Haupt, Conny Steiert, Laura Melchionda, Antoine Feignier, Karina von der Malsburg, Wolfgang Bildl, Jean-Baptiste van den Broucke, Catrin S. Müller, Thomas Becker, Martin van der Laan, Sven Dennerlein, Heike Rampelt, Bernd Fakler, Nils Wiedemann, Nikolaus Pfanner

## Abstract

Mitochondria function as cellular powerhouses and central hubs in metabolism, redox and stress reactions, signaling and apoptosis^1–5^. Defects of mitochondria lead to numerous human diseases^1,6–8^. The integration of mitochondrial proteins into complexes and networks is crucial for their function. Whereas the composition of the human mitochondrial proteome has been studied^8,9^, only limited information is available on the organization of the proteome into protein complexes and assemblies. Here we present a systematic mapping of the human mitochondrial complexome from HEK293T cells at super-resolution, resolving more than 7,000 abundance profile peaks of mitochondrial proteins. Proteins functioning in signaling, cell stress, protein biogenesis, turnover and membrane dynamics display particularly high complexities. High resolution and precise quantification enable discrimination between canonical constituents and non-stoichiometric regulatory interactors of the ATP synthase, major metabolite channels and import translocases. The complexome reveals membrane-spanning networks of protein insertase and morphology machinery, and co-assembly of protein import and export components at the major respiratory supercomplex, unraveling a multifunctional organization of mitochondrial machineries. This complexome represents a fully interactive resource for the systematic analysis of human mitochondrial machineries and interaction networks.

## Main Text

Nearly all human cells rely on the activity of mitochondria for the synthesis of large amounts of ATP to drive cellular processes. Although the role of mitochondria as cellular powerhouses has been known best, mitochondria perform a multitude of functions that are crucial for the proper development, growth and activity of human cells, including numerous metabolic pathways from synthesis of iron-sulfur clusters to amino acids, lipids and heme, signaling and stress responses, redox regulation, quality control and regulation of programmed cell death^1–5,10^. A large number of human diseases have been linked to mitochondrial defects and display a broad variety of phenotypes. More than 400 genes coding for proteins of the core mitochondrial proteome have been genetically associated with clinical phenotypes^1,6–9,11,12^.

Mitochondria have retained their own genetic system according to the endosymbiont hypothesis of mitochondrial origin from α-proteobacteria. The human mitochondrial genome codes for 13 hydrophobic proteins of the oxidative phosphorylation (OXPHOS) machinery comprising the respiratory chain and the F_1_F_o_-ATP synthase^6,13,14,15^, whereas the remaining 99% of mitochondrial proteins are encoded by nuclear genes and imported from the cytosol^16–18^. The mitochondrial membranes harbor elaborate systems for the transport of precursor proteins and their assembly into active machineries^17–21^. For the respiratory complexes I, III and IV, and the F_1_F_o_-ATP synthase, this involves the coordinated assembly of nuclear- and mitochondrial-encoded proteins^6,13,22–26^. Studies in various organisms revealed interactions between mitochondrial machineries of diverse functions, suggesting the formation of multifunctional networks^16,27–30^. The integration of different functions into coordinated networks is crucial for understanding the organization and cellular importance of mitochondria under physiological and pathophysiological conditions. However, the interaction networks of human mitochondria are only understood in part.

Complexome profiling uses mild separation of protein complexes coupled to mass spectrometric (MS) analysis, with native gel electrophoresis yielding the highest resolution^9,27,28^. Blue native electrophoresis-based mitochondrial complexomics started with 24 fractions (slices) per gel^31^, and most current studies use <120 slices per gel, with an average of ∼60 slices also for large multiomic studies^9,27,30,32–36^ (Fig. 1a). Although co-migration of proteins can be observed under these conditions, peaks with small molecular mass differences and subpopulations ‘buried’ under profile shoulders cannot be separated. The development of cryo-slicing blue native complexomics enabled use of the high blue native gel-resolving power in studies on rat and yeast mitochondria^28,37^. Here we report the highest-resolution complexome analysis (386 slices per gel) for HEK293T mitochondria (Fig. 1a). Combined with substantial improvement of mild mitochondrial lysis, blue native separation and quantitative MS analysis, the human mitochondrial complexome (hMitCOM) provides deep insight into the organization and interaction of mitochondrial protein complexes and supercomplexes of >900 different mitochondrial proteins, of which >400 are linked to human diseases.

**Fig. 1.**
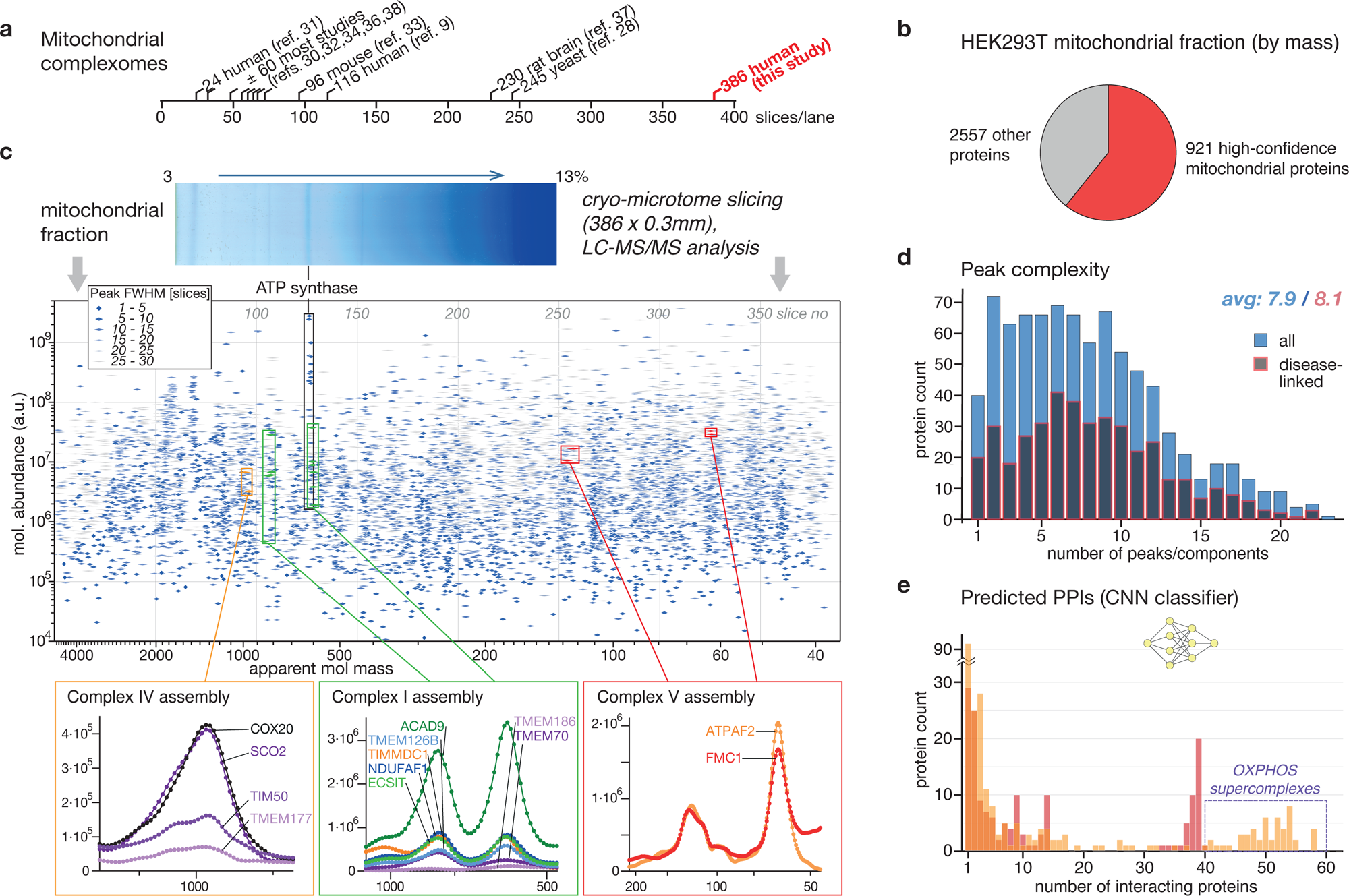
Complexome profiling of mitochondria. **a**, Overview of blue native complexome profiling studies of mitochondria by their sampling resolution, ranging from 24 to 116 slices per lane by conventional slicing, and up to 386 by cryo-microtome-assisted slicing. **b,** Composition of the HEK293T mitochondrial fraction by the summed abundances (i.e. profile integrals) of the identified proteins. About 60% of the protein mass represent 921 high-confidence mitochondrial proteins (Supplementary Tables 1 and 2), whereas the remaining portion is shared by 2557 proteins. **c,** Top: preparative 3-13% blue native gel lane of HEK293T mitochondrial fraction (lysed with digitonin) subjected to cryo-microtome slicing into 386 samples. Middle: summary plot visualizing individual peak components (quadrangular symbols) by their molecular (mol.) abundance (y-axis), maximum slice index (upper x-axis, grey) or apparent molecular mass (lower x-axis, black) as determined by unsupervised multi-gaussian fits to the respective protein abundance profiles of the HEK293T-MitCOM core proteins. Symbols indicate ranges of the full-width at half-maximal height of the fitted components. The prominent band in the blue native-gel aligns with the main peak components of F_1_F_o_-ATP synthase (complex V, boxed black). Bottom: insets showing zooms of abundance-mass profiles of OXPHOS complex assembly factors forming novel assemblies. Boxes in the peak component plot highlight the respective fitted profile components. **d,** Histogram depicting the peak complexity distribution based on the number of fitted components/protein profile. Blue bars show the total protein counts per bin, dark bars framed red indicate the counts of disease-linked proteins per bin (making up 48 % of HEK293T-MitCOM core). Peak component averages are given for both sets of proteins. **e,** Assembly complexity assessed by counting the number of profiles predicted to represent interacting proteins (predicted protein-protein interactions (PPIs), convoluted neuronal network classifier (CNN) score >0.5; Methods and Extended Data Figs. 2 and 3). A total of 1765 mitochondrial protein pairs were predicted as positive. The boxed region (>40 interactors per protein) reflects OXPHOS subunit integration into supercomplexes.

### Super-resolution complexome of human mitochondria

The mitochondrial fraction was prepared from HEK293T cells and its complexome analyzed by blue native gel separation and quantitative MS (Fig. 1b,c). To maintain mitochondrial protein network interactions and to achieve a high complexome resolution, we focused on the following aspects: (1) Mitochondria were lysed under very mild conditions using the non-ionic detergent digitonin at a 2.5:1 ratio to mitochondrial protein, compared to typical digitonin/protein ratios of 4:1 to 9:1 (average 6:1)^9,30,34,38^. (2) The blue native gradient gel separation was optimized for preparative-scale extended separation range for the expected higher molecular mass assemblies (Fig. 1c). (3) Cryo-slicing of the blue native gel yielded 386 slices of 0.3 mm width. Since the sharpest-focusing protein bands exhibited a half-width of 1.2-1.5 mm, sampling covered each abundance profile peak by at least four slices (Fig. 1c and Extended Data Fig. 1a). (4) Data processing, in particular run-to-run normalization and noise removal, were improved. (5) Manual evaluation of protein abundance profiles was complemented by component-detection and AI-empowered prediction of preferred protein assemblies.

Of 3,478 proteins identified in the mitochondrial preparation, 921 proteins were assigned as high-confidence mitochondrial proteins and yielded hMitCOM profiles (Fig. 1b and Supplementary Tables 1 and 2, www.complexomics.org). The profiles cover a molecular mass range of 40 kDa to 4,700 kDa and a protein abundance range of more than five orders of magnitude (Fig. 1c and Extended Data Fig. 1a,b). The exquisite resolution and precision of the profiles permit a systematic analysis of human mitochondrial protein complexes, supercomplexes and assembly intermediates.

851 protein profiles were subjected to an automated component analysis consisting of a peak detection and multi-gaussian fitting procedure. An average of 7.9 profile peak components per protein revealed a high complexity of the human mitochondrial proteome organization (Fig. 1d). Furthermore, we trained a convoluted neuronal network classifier (CNN) to predict protein-protein interactions (PPIs) based on co-migration patterns of their respective abundance profiles (Fig. 1e and Extended Data Figs. 2 and 3). Since we applied rather stringent parameters for these evaluations, the determined peak components and predicted PPIs likely even underestimate the complexity of the human mitochondrial proteome. The profiles can be analyzed in an openly accessibly interactive profile viewer, which includes a correlation analysis feature to search for protein interaction candidates and complexes (www.complexomics.org).

### Complexity of human mitochondria, functional classification and disease association

We analyzed the complexity of human mitochondrial proteins according to their functional classification (Fig. 2a,b). The three categories signaling & cell stress, protein biogenesis & turnover, and mitochondrial morphology & membrane dynamics showed the highest complexity compared to the average complexity of 7.9 peaks per hMitCOM protein. The complexity pattern across functional protein categories is strikingly different between human mitochondria and mitochondria from the lower eukaryote baker’s yeast. In yeast, oxidative phosphorylation shows the highest complexity, whereas the three categories with highest complexity in human mitochondria are only in the average range or even below in yeast mitochondria^28^. Thus, human mitochondria display particularly high complexities in three functional categories that are important for the integration of mitochondria into the cellular organization: signaling, protein biogenesis and membrane dynamics. Abundant housekeeping categories such as metabolism and mitochondrial gene expression showed a below average complexity in human mitochondria (Fig. 2b).

**Fig. 2.**
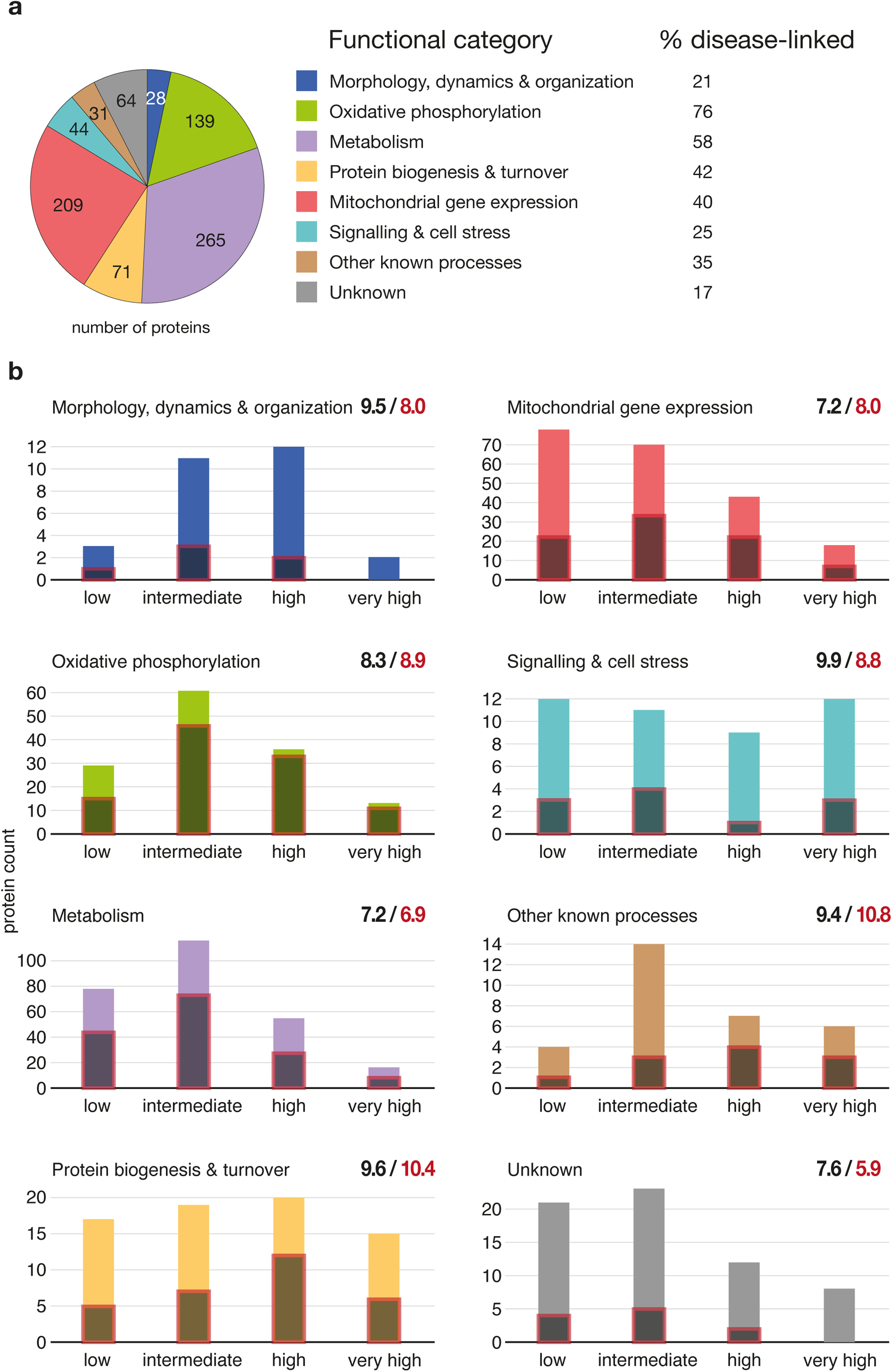
Functional classification, disease-association and complexity of hMitCOM proteins. **a**, Representation of hMitCOM core proteins in eight functional categories defined by their database (UniProt)-assigned primary molecular function. The percentage of disease-linked proteins are shown per category. **b,** Histograms showing hMitCOM core protein profile complexity binned in categories “low” (1-4), “intermediate” (5-9), “high” (10-14) and very high (>15 peak components per profile) for the indicated functional categories. Superimposed bars framed red represent disease-linked proteins. Component count averages are indicated by numbers (hMitCOM core proteins black, fraction of disease-linked proteins red).

A systematic survey of the disease association of hMitCOM revealed that 48% of the proteins analyzed were encoded by genes linked to human diseases^12,39^ (Figs. 1d and 2a,b, Supplementary Tables 2 and 3 and Methods). Whereas the long established functional mitochondrial categories oxidative phosphorylation, metabolism, and mitochondrial gene expression contained a considerable fraction of disease-linked genes as expected, the category protein biogenesis & turnover not only included a considerable fraction but also a remarkably high complexity of over 10 for disease-linked proteins (Fig. 2b). This underscores the growing evidence for an important role of protein biogenesis and turnover for human mitochondrial pathophysiology^5,6,16,40^. On the other hand, the category ‘proteins with unknown function’ contained the lowest fraction and the lowest complexity of disease-linked genes. The detailed functional classifications and disease-associations of the hMitCOM proteins are outlined in the Supplementary Tables 2 and 3. In summary, nearly half of hMitCOM genes are disease-linked and include many high to very high complexity proteins (Figs. 1d and 2b).

### Separation of stoichiometric and regulatory components of TOM-VDAC and mitochondrial ATP synthase

We asked if the quantitative high-resolution mapping by hMitCOM made it possible to discriminate between canonical (stoichiometric) components and non-stoichiometric regulatory partners of mitochondrial protein complexes. The protein translocase of the outer membrane (TOM) has been reported to interact with the mitochondrial metabolite channel porin, also termed voltage-dependent anion channel (VDAC)^37,41–44^. Different roles have been ascribed to this interaction from transient interactions of TOM and VDAC to binding of the unassembled TOM22 subunit by VDAC for regulation of protein import. A cryo-electron microscopy study of the human TOM complex upon depolarization of mitochondria and preprotein arrest unexpectedly showed a stoichiometric array of mature TOM and VDAC complexes^45^. However, it has been unknown whether stoichiometric TOM-VDAC arrays are a common feature of the mitochondrial outer membrane or only observed under special conditions^45^. Since hMitCOM displayed quantitative profiles from HEK293T cells grown under standard conditions, we directly compared TOM and VDAC complexes and observed a perfect co-migration of the TOM core components TOM40 and TOM22 with VDAC1, VDAC2 and VDAC3 (Fig. 3a,b and Extended Data Fig. 4a,b). For comparison, the hMitCOM profile of the pro-apoptotic protein BAK, a well-established interaction partner of VDAC2 (refs.^46–50^), only partially overlapped with a shifted maximum compared to the VDAC2 profile (Fig. 3b). Direct quantitative comparison demonstrated that mitochondrial BAK was more than an order of magnitude less abundant than VDAC2 (Extended Data Fig. 4c). Under these non-apoptotic conditions, BAK is not a stoichiometric partner of VDAC2, but rather associates with a small fraction of VDAC complexes, leading to a ‘buried’ subpopulation of molecular mass-shifted VDAC2-BAK complexes. In contrast, quantification of the TOM/VDAC peaks revealed that TOM40 was in a stoichiometric range with regard to VDAC complexes (Extended Data Fig. 4a,b). Together with the perfect overlap of the major TOM and VDAC peak (Fig. 3b) and TOM-VDAC interactions in pull-down and structural studies^37,41–45^, these data indicate that stoichiometric TOM-VDAC assemblies are present under standard growth conditions without special treatment of human mitochondria.

Omic studies reported an association of the protein C15orf61 with the F_1_F_o_-ATP synthase and the mitochondrial inner membrane^9,30,51^. The hMitCOM dataset indeed revealed a strong profile correlation between C15orf61 and established subunits of the ATP synthase (complex V) (Fig. 3c; Pearson correlation coefficient up to 0.97). Tagged C15orf61 specifically co-purified ATP synthase subunits from digitonin-lysed mitochondria, but not subunits from respiratory complexes I, II, III or IV (Fig. 3d and Extended Data Fig. 5a). Blue native electrophoresis showed that the assembled ATP synthase complex was associated with C15orf61 (Fig. 3e), and C15orf61 was not extracted from the membranes at alkaline pH (Fig. 3f), indicating that C15orf61 is a membrane protein interacting with the assembled ATP synthase. Upon knockout of C15orf61, the levels of ATP synthase subunits and further mitochondrial proteins remained largely unaffected^30^, only the levels of the regulatory inhibitor protein ATP5IF1 were moderately increased (Extended Data Fig. 5b). The abundance of C15orf61 was reported to be negatively correlated with respiratory conductance^52^, and in C15orf61 knockout mitochondria, increased levels of ATP synthase dimers and oligomers were observed at the expense of monomers, indicating that C15orf61 negatively regulated ATP synthase multimerization^30^. It was concluded that C15orf61 represented a new stoichiometric subunit of the mitochondrial ATP synthase^30^. Quantitative mapping of hMitCOM profiles, however, demonstrated that C15orf61 was more than an order of magnitude less abundant than established ATP synthase subunits (Fig. 3g). The high-resolution hMitCOM mapping revealed that C15orf61 was not part of the major monomeric ATP synthase peak of 600 kDa, but shifted to a higher molecular mass peak (Fig. 3h), fully supporting that C15orf61 was specifically associated with a small fraction of ATP synthase complexes. We conclude that C15orf61 is a non-stoichiometric interactor of ATP synthase complexes, consistent with regulatory roles in the function and organization of the mitochondrial ATP synthase^30,52^.

**Fig. 3.**
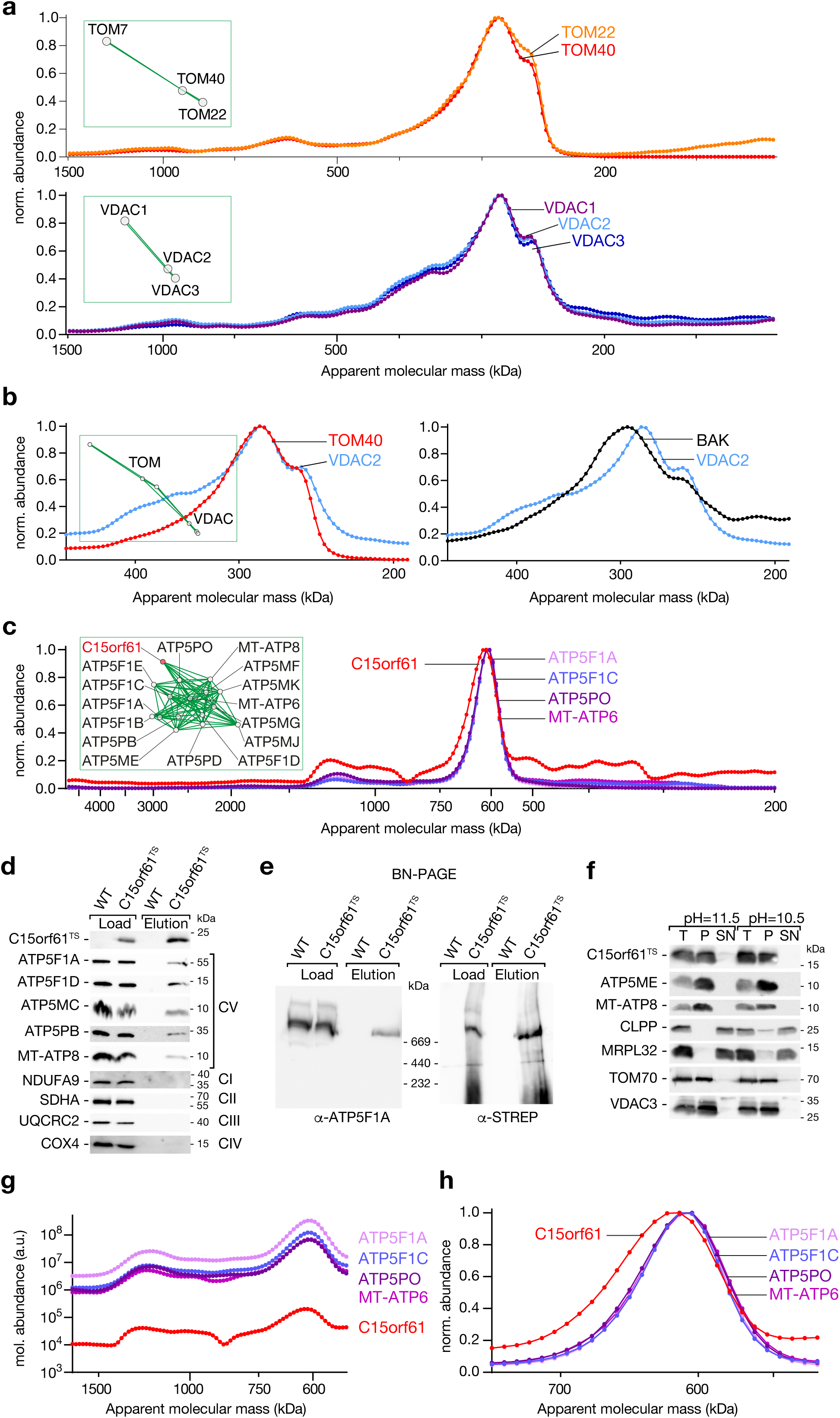
Separation of stoichiometric subunits and non-stoichiometric interactors by quantitative profiling. **a**, Normalized (norm.) abundance-mass profiles of human TOM40 and TOM22 (top) and VDAC1, VDAC2 and VDAC3 (bottom). Insets, predicted protein-protein interactions (PPIs) based on convoluted neuronal network classifier (CNN). **b**, Comparison of normalized abundance-mass profiles of VDAC2 with TOM40 or BAK, respectively. Inset, CNN-based predicted PPIs. **c**, Normalized abundance-mass profiles of C15orf61 and F_1_F_o_-ATP synthase subunits. Inset, CNN-based predicted PPIs. **d**,**e**, Mitochondria from HEK293T cells expressing C15orf61-TwinStrep were solubilized in lysis buffer containing 1% digitonin and subjected to Twin-Strep affinity purification. Co-purified complexes were eluted with Biotin-containing elution buffer and analysed by SDS-PAGE (**d**) and blue native-PAGE (**e**). Load 5%, eluate 100%. WT, wild-type. **f**, Mitochondria were subjected to carbonate treatment and analyzed by SDS-PAGE. **g**, Molecular (mol.) abundance-mass profiles of C15orf61 and F_1_F_o_-ATP synthase subunits. **h**, Magnified view of **c**, showing the shift of the C15orf61 peak in comparison to the F_1_F_o_-ATP synthase monomer peak.

The precision of hMitCOM in both molecular mass separation and absolute quantification of individual protein peaks thus enables efficient separation of stoichiometric subunits from non-stoichiometric interaction partners of protein complexes.

### Organization of MICOS-SAM supercomplexes spanning two membranes

The mitochondrial contact site and cristae organizing system (MICOS) of the inner membrane stabilizes crista junctions between inner boundary and cristae membranes and forms contact sites to the outer membrane^53–62^. Classical blue native gel separation and immunoblotting yielded three main MICOS bands in human mitochondria, similar to intermediate-resolution complexomics^9,38^ (Fig. 4a). hMitCOM resolved a ∼560 kDa MICOS core complex and a series of larger complexes (Fig. 4b). Human MICOS consists of seven different subunits with MIC60 and MIC10 as the major constituents; it has been assumed that all seven subunits together form the MICOS core complex^38,63,64^. However, MitCOM revealed that MIC27 did not co-migrate with the MICOS core complex in contrast to all other MICOS subunits (Fig. 4c). A quantitative peak analysis showed that MIC27 was considerably less abundant than the homologous subunit MIC26. The main peak of MIC27 was shifted compared to the MICOS core peak and aligned with a MICOS core shoulder (Extended Data Fig. 6a, asterisk). We conclude that MIC27 is not a stoichiometric MICOS core component, it associates only with a fraction of the core complexes and may play a regulatory role^65^.

**Fig. 4.**
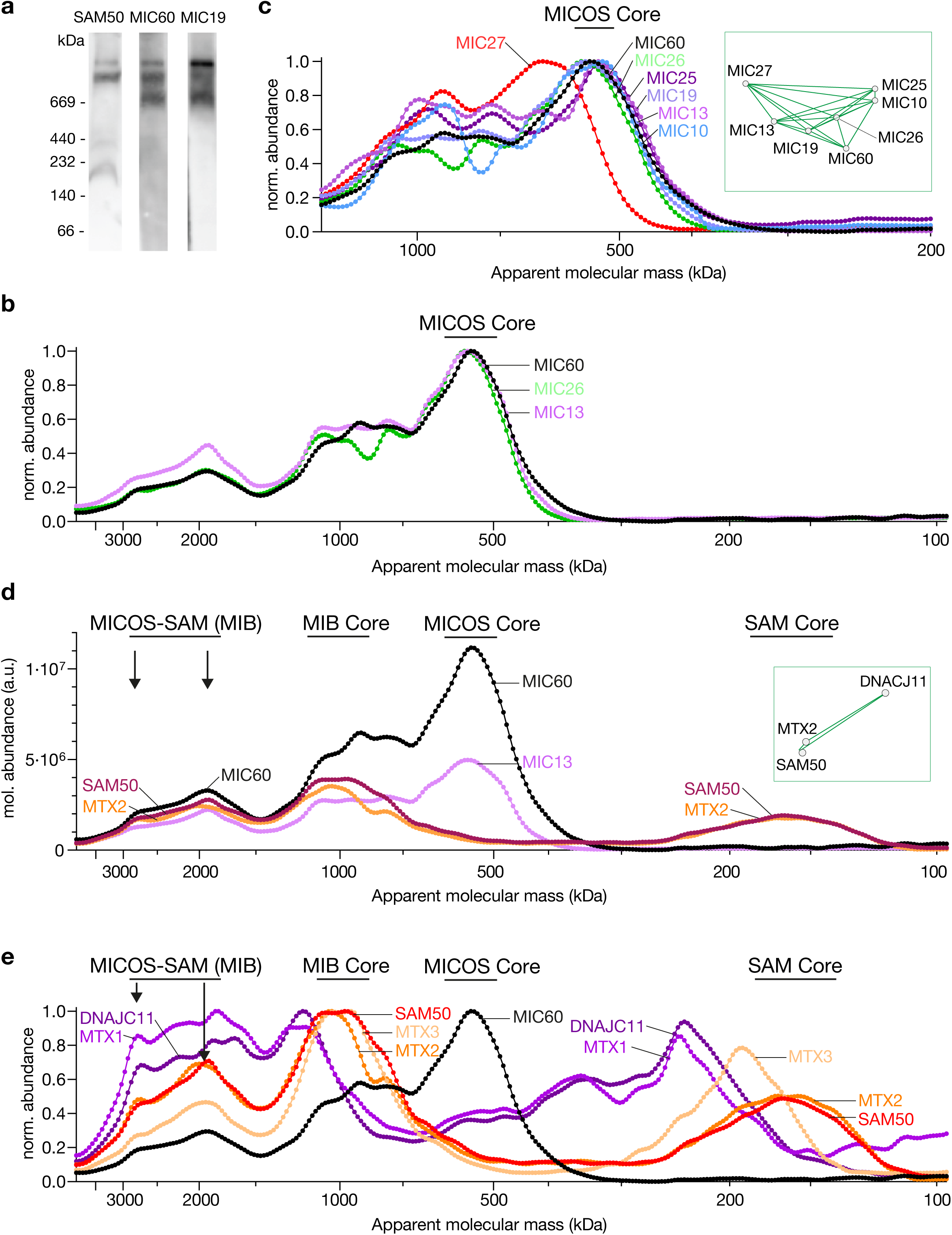
Organization of MICOS and MICOS-SAM supercomplexes. **a**, SAM50, MIC60 and MIC19 complexes were visualized from isolated HEK293T mitochondria after solubilization in digitonin, separation by blue native-PAGE and detection by immunodecoration. **b**,**c**, Normalized (norm.) abundance-mass profiles of MICOS subunits. Inset, CNN-based predicted PPIs. **d**, Molecular (mol.) abundance-mass profiles of MICOS and SAM subunits and supercomplexes. Inset, CNN-based predicted PPIs. **e**, Migration of the additional SAM subunits (MTX1, MTX2, MTX3 and DNAJC11) in comparison to SAM50 and MIC60 in normalized abundance-mass profiles. MICOS-SAM (MIB) supercomplexes, MIB Core, MICOS Core and SAM Core are indicated.

Human MICOS associates with the sorting and assembly machinery (SAM) complex of the outer membrane, forming two-membrane-spanning MICOS-SAM supercomplexes migrating in the range of 1-3 MDa, also termed mitochondrial intermembrane space bridging (MIB) complex^38,56,57,63,66–69^ (Fig. 4a,d,e and Extended Data Fig. 6a,b). It has been assumed that SAM50 and Metaxins 2 and 3 (MTX2, MTX3) together form the SAM core complex^38^ (Extended Data Fig. 7). However, MTX3 was not present in the ∼160 kDa SAM core complex containing SAM50 and MTX2, but co-migrated with SAM50/MTX2 and MICOS in the 1 MDa MIB core complex and the 2 MDa supercomplex (Fig. 4d,e). All known SAM components, including MTX1 and DNAJC11 (refs.^38,70^), co-migrated in the full MICOS-SAM supercomplex of 2.8 MDa. We conclude that neither MICOS core nor SAM core contain all subunits that have been assigned to them. However, in the large two-membrane-spanning MICOS-SAM supercomplex, all components are present. Thus, human MICOS and SAM stabilize each other across two membranes, providing an explanation for the puzzling observation^71^ that substrates (β-barrel precursors) of human SAM were found in high-molecular mass intermediates during their assembly, rather than in small SAM complexes^67,71,72^. Human SAM is only complete in the MICOS-SAM supercomplexes and thus the presence of assembly intermediates fully agrees with the view that the supercomplexes represent active forms.

Quantitative comparison revealed that the SAM core complex was of much lower abundance than the MICOS core complex, however, their abundance came close in the 1 MDa MIB core and was nearly stoichiometric in the full MICOS-SAM supercomplexes (Fig. 4d and Extended Data Fig. 6b). Thus, stoichiometric arrays of MICOS and SAM form two-membrane-spanning supercomplexes and provide the framework for the organization of inner membrane crista junctions underneath the mitochondrial outer membrane and, potentially, targeted/local β-barrel protein biogenesis^38,56,57,66–69,71,73^.

### Protein import and export machineries at respiratory supercomplex

The mitochondrial inner membrane contains three protein translocases. The presequence translocase (TIM23 complex) and the carrier translocase (TIM22 complex) mediate the import of nuclear-encoded precursor proteins, whereas OXA1L (oxidase assembly) functions as export translocase and inserts mitochondrial-encoded proteins and some matrix-sorted nuclear-encoded proteins into the inner membrane^17–21,74–78^. The three translocases migrate at different positions in the lower molecular mass range of hMitCOM (Fig. 5a). The presequence translocase subunit TIM21, however, showed a peculiar behavior in the high molecular mass range (Fig. 5a). TIM21 has been known to promote the transfer of precursor proteins from the presequence translocase to assembly intermediates of respiratory chain complexes that migrate below the 1 MDa range, particularly the mitochondrial translation regulation assembly intermediate of cytochrome c oxidase (MITRAC) running at ∼200 kDa^79,80^. hMitCOM revealed a significant TIM21 peak at 1.8 MDa; for comparison, the major respiratory supercomplex containing complexes I, III and IV migrated at ∼1.5 MDa (Fig. 5a,b). Strikingly, the export translocase OXA1L also displayed a peak at 1.8 MDa (Fig. 5a). OXA1L interacts with early assembly intermediates of respiratory chain complexes^74,81^ distinct from TIM21-containing intermediate complexes. Neither an interaction of TIM21 and OXA1L nor their presence in complexes larger than the mature respiratory supercomplex have been reported. We thus performed affinity purification from digitonin-lysed mitochondria using either tagged TIM21 or tagged OXA1L. Under both conditions, we observed a co-purification of TIM21 and OXA1L (Fig. 5c). The 1.8 MDa TIM21/OXA1L peak migrated in a small shoulder of the respiratory supercomplex (Fig. 5a) and indeed we observed a co-purification of OXA1L, TIM21 and the complex I subunit NDUFA9 (Fig. 5c). A detailed inspection demonstrated the precise co-migration of OXA1L and TIM21 at 1.8 MDa at a shoulder of the respiratory supercomplex (Fig. 5d), which we termed Mitochondrial TIM-OXA-Respiratory chain (MiTOR) peak. The absolute quantification showed the considerably lower abundance of OXA1L and TIM21 compared to the respiratory supercomplex, explaining why MiTOR did not lead to a second major peak of the supercomplex, but was visible as a shoulder in the profile (Fig. 5d). We conclude that a fraction of the major respiratory supercomplexes of the mitochondrial inner membrane are associated with protein import and export machineries simultaneously.

**Fig. 5.**
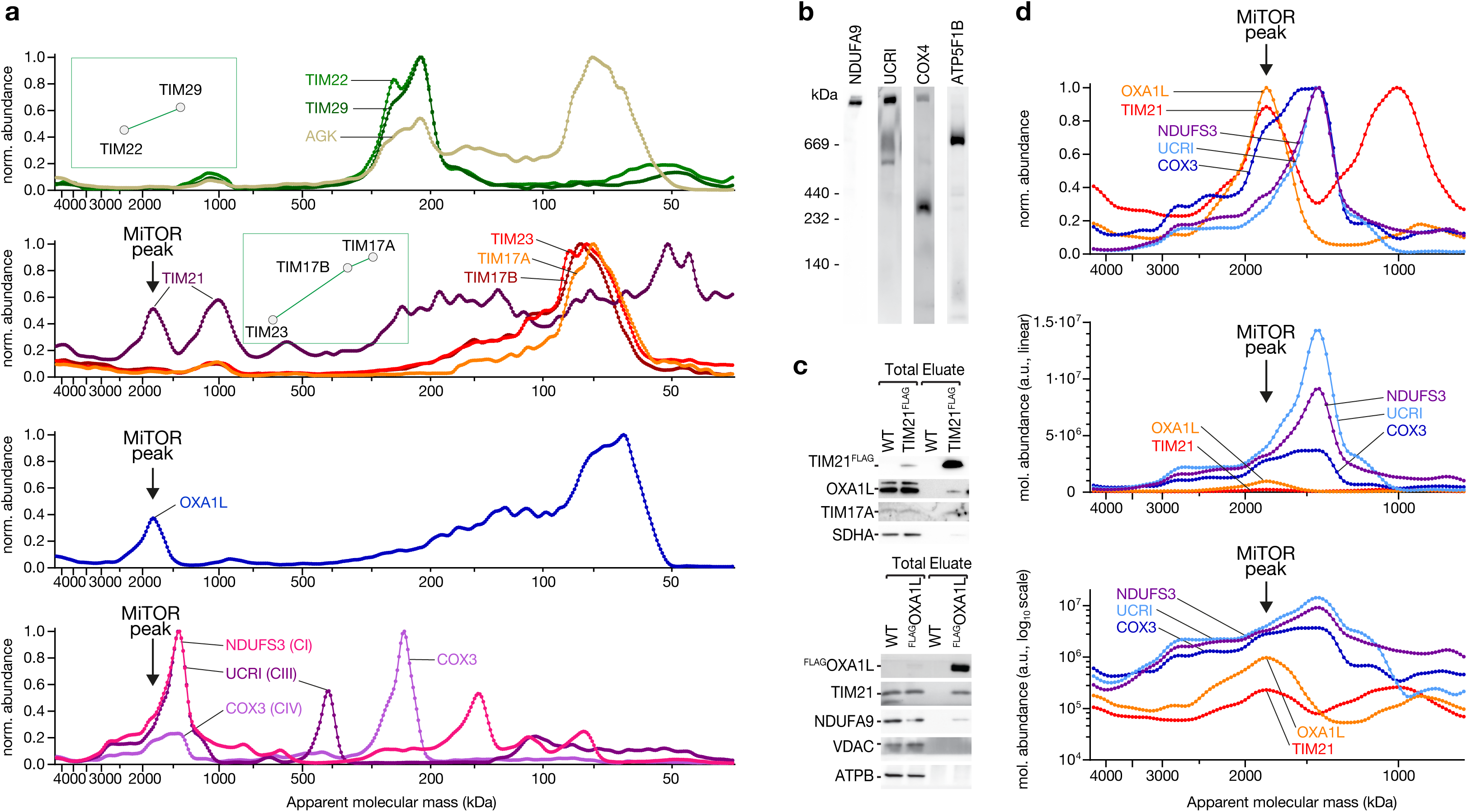
MiTOR import and export machineries at respiratory supercomplex. **a**, Normalized (norm.) abundance-mass profiles of carrier import translocase (TIM22) subunits (top), presequence import translocase (TIM23) subunits (second diagram), OXA1L export translocase (third diagram) and respiratory complexes (C) I, III and IV (bottom). Insets, CNN-based predicted PPIs. **b**, Isolated mitochondria from HEK293T cells were solubilized in digitonin and subjected to blue native-PAGE. OXPHOS complexes (CI–NDUFA9, CIII–UCRI/Rieske, CIV–COX4, and CV–ATP5F1B) were detected by immunodecoration and chemiluminescence. **c**, Immunoprecipitation from control (WT, wild-type), TIM21^FLAG^ or ^FLAG^OXA1L containing mitochondria. Solubilized mitochondria were subjected to anti-FLAG agarose and bound proteins eluted by FLAG-peptide. Samples were subjected to SDS-PAGE and immunodecoration. Total 0.75%, eluate 100%. **d**, Magnified areas and comparison of the co-migration of TIM21, OXA1L and respiratory subunits in normalized and molecular (mol.) abundance-mass profiles.

The quantitative dissection of mitochondrial protein distribution over the entire molecular mass range combined with biochemical analysis thus reveals a megadalton-organization of mitochondrial machineries that had only been studied in lower molecular mass entities before.

## Conclusions

The HEK293T human mitochondrial complexome dissects more than 7,000 distinct protein peaks, providing a resource with an enormous complexity of mitochondrial protein assemblies, supercomplexes, intermediates and networks. Native mitochondrial protein complexes solubilized under mild conditions were resolved without tagging or other modifications. The extraordinary resolution enables the separation of protein peaks and shoulders with small molecular mass differences at sub-stoichiometric level. Together with the direct quantitative comparison of individual peak components, functional classification and biochemical analysis, the complexome provides deep insight into the organization of mitochondrial protein networks.

The following examples illustrate how hMitCOM can be used to systematically unravel protein networks of human mitochondria. (1) hMitCOM enables direct separation of stoichiometric subunits and non-stoichiometric interactors of protein complexes. We elucidate that the protein translocator TOM40 is a stoichiometric partner of VDAC metabolite channels, whereas C15orf61 is a non-stoichiometric interaction partner of the mitochondrial ATP synthase in contrast to the current assumption^30^. (2) The inner membrane morphology machinery MICOS and the outer membrane protein insertase SAM form stoichiometric arrays of nearly 3 MDa, containing all known MICOS and SAM subunits. In contrast, neither the MICOS core complex nor the SAM core complex contain all assigned subunits. Thus, human MICOS and SAM stabilize each other in two-membrane-spanning megacomplexes, explaining the puzzling observation that the substrates of human SAM were not found in association with SAM core, but in high-molecular mass intermediates during their assembly^71,72^. (3) The import component TIM21 and the export translocase OXA1L function in respiratory chain biogenesis that involves assembly of nuclear-encoded proteins and mitochondrial-encoded proteins^14,15,79,81^. TIM21 and OXA1L have been observed in separate assembly intermediates in the lower molecular mass range and a physical association has not been reported. hMitCOM discovered the MiTOR peak, which is larger than the major respiratory supercomplex and contains both TIM21 and OXA1L in precise co-migration. Together with biochemical affinity purification and quantitative analysis of a supercomplex shoulder, we show that a fraction of the major respiratory supercomplexes are associated with protein import and export components simultaneously. The presence of import and export machineries at mature respiratory supercomplexes opens a new view on the mechanisms of continuous remodeling of the mitochondrial respiratory chain.

hMitCOM is freely available as an interactive and searchable resource for the scientific community at www.complexomics.org. Since nearly half of the genes coding for hMitCOM proteins have been linked to human diseases (Fig. 2 and Supplementary Tables 2 and 3), we anticipate that hMitCOM will not only provide systematic information about mitochondrial protein assemblies and their composition, but also insight into the networks and assembly processes of the large number of disease-linked gene products.

## Methods

### Cell growth for cryo-slicing analysis

HEK293T cell were cultured in a humidified incubator with 5% CO_2_ at 37°C. Cells were maintained in high glucose DMEM (Gibco) containing 10% fetal bovine serum (FBS, Anprotec), 1% penicillin-streptomycin (Gibco) and 2 mM L-Glutamine (Gibco). Cells were grown to approximately 80% confluency and subsequently passaged.

### Cloning

The target genes were cloned into the pcDNA3.1 (+) mammalian expression vector using Gibson assembly. Briefly, the insert and vector fragments were generated with overlapping sequences via PCR using the KOD Hot Start Polymerase (Sigma-Aldrich). The fragments were then assembled in a one-step assembly reaction using the Gibson Assembly Cloning Kit (NEB). The final assembled plasmid was transformed into chemical competent *E. coli*, isolated using the Qiagen Mini Prep Kit (Qiagen).

### Transfection for C15orf61 analysis

HEK293T cells were transfected, using FuGENE®6 transfection reagent (Promega) according to the manufacturers protocol. In summary, HEK293T cell were expanded in a 150 mm dish to approximately 70% confluency. Prewarmed media and DNA were mixed and the transfection reagent was added in a 3:1 FuGENE to DNA ratio. The transfection mixture was incubated at room temperature for 15 min, added to cells and incubated overnight. On the following day, the media was replaced and 48 hrs post transfection, the cells were split and plated in selective medium, containing 400 µg/ml Zeocin (Invitrogen). One plate of cells was used to assess transfection efficiency and the second plate was used for the generation of a stable cell line through continuous growth in selective media over two weeks until distinct colonies formed. Clones were obtained by sorting single cells into individual wells of a 96 well plate, followed by clonal expansion.

### CRISPR/CAS9 knockout

The oligonucleotides targeting the C15orf61 target sequence were designed using the online bioinformatics tool CHOPCHOP^82^ (http://chopchop.cbu.uib.no). The oligonucleotides were phosphorylated, annealed and ligated into the BbsI-digested pSpCas9-(BB)-2A-GFP vector as published before^83^. After transfection of the CRISPR/CAS9 plasmid, the cells were washed in DPBS (PAN Biotech) and resuspend in 5 ml DPBS containing 2% FBS. Cells were sorted for GFP expression by flow cytometry and single-cell seeded into 96 well plates. The resulting single-cell clones were subsequently expanded and screened for the knockout of the target gene. For this, the gene of interest was amplified by PCR, cloned into a Zero Blunt PCR Cloning Kit vector, transformed into TOP10 *E. coli*. Subsequently, plasmids of individual colonies were isolated and sequenced. Plasmids and primers are listed in Supplementary Table 4.

### Mitochondrial preparation

Mitochondria were isolated as described previously^84^. Briefly, cells were harvested in 10 ml PBS, centrifuged at 800 x g for 5 min and were frozen at −80°C to induce cell lysis. The cell pellet was resuspended in 10 pellet volumes of buffer A (83 mM sucrose, 10 mM HEPES, pH 7.2) and transferred to a glass homogenizer. Cells were homogenized and an equal volume of buffer B (250 mM sucrose, 30 mM HEPES, pH 7.2) was added to the homogenate. To remove cell debris, the homogenate was centrifuged at 1,000 x g for 5 min. The supernatant was moved to a fresh microcentrifuge tube and mitochondria were pelleted at 12,000 x g for 2 min at 4°C. The mitochondrial pellet was resuspended in buffer C (320 mM sucrose, 1 mM EDTA, 10 mM Tris, pH 7.4). Protein concentration was determined using Bradford assay. Mitochondria were aliquoted and snap frozen in liquid nitrogen to be stored at −80°C.

### Blue native gel electrophoresis for cryo-slicing analysis

3 to 13% non-denaturing blue native polyacrylamide gels were cast to separate proteins under native conditions^85^. For the cryo-microtome slicing, the gels were cast continuously without a stacking layer, using 1.5 mm spacers and a comb that formed two 3.2 cm wide sample wells. Other gels were prepared with a stacking and separating layer, using 1 mm spacers and standard 15 well combs. TEMED and ammonium persulfate were freshly added to gel solution A (12.25% acrylamide, 0.74% bis-acrylamide, 67 mM aminocaproic acid, 50 mM Bis-Tris, 20% glycerol [v/v]) and gel solution B (2.92% acrylamide, 0.08% bis-acrylamide, 67 mM aminocaproic acid, 50 mM Bis-Tris). Both solutions were mixed and poured using a peristaltic pump. The gels were allowed to polymerize overnight and subsequently run in a cooled Hoefer vertical protein electrophoresis unit with a cathode (50 mM Tricine, 50 mM Bis-Tis, 0.02% [w/v] Coomassie G, pH 7.0) and anode (50 mM Bis-Tris, pH 7.0) buffer system. For the cryo-microtome gel, 1 mg of human HEK293T mitochondria were solubilized with digitonin (Matrix Bioscience) at a 2.5:1 [w/w] digitonin-to-protein ratio in solubilization buffer (20 mM Tris-HCl, pH 7.4, 50 mM NaCl, 10% [v/v] glycerol) to a final concentration of 4 µg/µl and centrifuged at 18,000 x g to remove insoluble material. For all samples, 10x loading dye (100 mM Bis-Tris, 500 mM aminocaproic acid, 5% [w/v] Coomassie G, pH 7.0) was added to the cleared supernatant and the samples loaded onto the gel. Electrophoresis was performed at 4°C overnight at a maximal current of 6 mA and/or a maximal voltage of 70 V. On the next day the electrophoresis was continued at a maximal voltage of 600 V.

### Cryo-slicing blue native mass spectrometry (csBN-MS)

The blue native gel lane was excised, fixed in 15% acetic acid / 30% ethanol (v/v), then embedded in tissue embedding medium (LEICA) and frozen at −20°C. Cryo-microtome slicing (Leica CM 1950, 0.3 mm step size) was carried out as described^86^ yielding 386 slice samples in total. In-gel tryptic digestion (sequencing grade modified trypsin; Promega GmbH, Germany) was performed in filtration plates (96-well format, Porvair Science) after extensive washing to remove residual polymers. Extracted peptides were dissolved in 20 µl 0.5% [v/v] trifluoroacetic acid, and 5 µl aliquots were loaded with 0.05% [v/v] trifluoroacetic acid onto a µPAC trapping column (PharmaFluidics / Thermo Fisher Scientific) via the autosampler of a split-free UltiMate 3000 RSLCnano HPLC (Thermo Fisher Scientific). Bound peptides were eluted and separated on a 50 cm µPAC C18 reversed phase analytical column (Gen1; PharmaFluidics / Thermo Fisher Scientific) with a FS360-20-10-N-5-105CT emitter (shortened to 5 cm; New Objective) using the following aqueous-organic elution gradient (eluent A: 0.1% [v/v] formic acid and eluent B: 0.1% [v/v] formic acid in 80% [v/v] acetonitrile): 5 min 3% ‘B’ (equilibration), 90 min from 3% to 30% ‘B’ and 15 min from 30% to 40% ‘B’ and 5 min from 40% to 50% ‘B’ and 5 min from 50% to 99% ‘B’ (four-step gradient), 5 min 99% ‘B’ (washing), 5 min from 99% to 3% ‘B’ and 15 min 3% ‘B’ (flow rate 300 nl/min). Measurements were performed on an Orbitrap Exploris 480 mass spectrometer with a Nanospray Flex NG ion source (both Thermo Fisher Scientific) using the following settings: spray voltage 2.3 kV, ion transfer tube temperature 275°C, advanced peak determination, cycle time 2.5 s, master scan orbitrap resolution 240,000, scan range 370-1,700 m/z, RF lens 40%, normalized AGC target 300%, maximum injection time 200 ms, profile data type, positive polarity, monoisotopic precursor selection (peptide) filter, intensity threshold 10,000, include charge states 2-4, skip undetermined charge states, dynamic exclusion duration 45 s with mass tolerance ±10 ppm; data-dependent MS/MS settings: isolation window 0.8 m/z, normalized HCD collision energy 30%, orbitrap resolution 15,000, first mass 100 m/z, normalized AGC target 75%, maximum injection time 200 ms, centroid data type.

### Protein identification and annotation

LC-MS/MS data files were recalibrated with mzRecal v1.1.4 (https://github.com/524D/mzrecal) and searched against the UniProt human reference proteome (release 2024_06) and GPM cRAP contaminants database with PEAKS Online 12 (Bioinformatics Solutions, Inc.) and the following settings: automatic mass correction, 5 ppm precursor and 0.02 Da fragment mass tolerance, up to 3 missed tryptic cleavage sites and up to 3 variable modifications per peptide (any of Acetylation (Protein N-term), Carbamidomethylation (C), Deamidation (NQ), Formylation, Oxidation (M), Pyro-glu from E, Pyro-glu from Q). In total, 4,625 protein groups with 105,104 unique peptidoforms were identified at 1% protein group FDR. Exogenous contaminants (e.g. keratins, trypsin, IgG chains) or proteins identified in less than three slices or those providing insufficient peptide information for determination of an abundance-mass profile (see below) were not considered further.

Of the remaining 3478 proteins (HEK293T complexome dataset), 921 were assigned high-confidence mitochondrial proteins (constituting hMitCOM) based on UniProt database, Human Protein Atlas (HPA, www.proteinatlas.org), Human MitoCarta3.0 (ref.^87^) and a landmark proteomic study (ref.^9^) as annotation sources as well as manual curation using literature. Additional features of hMitCOM proteins (gene name, protein name, number of amino acids, predicted molecular weight, transmembrane domains, lipid modifications, GO term assignments) were retrieved from UniProt (release 2025_03) (Supplementary Table 2 - hMitCOM).

851 hMitCOM core proteins exhibiting high quality profiles (as manually assessed based on representation by quantifiable peptides, protein abundance over detection threshold, profile continuity and noise of profile peaks) were selected for detailed systemic analyses, i.e. peak profile complexity (see below), functional classification by manual evaluation of GO term annotations and literature, or linkage to disease / phenotypes (see below) (Supplementary Table 2 - Evaluation). Information on genetic diseases and phenotypes linked to individual hMitCOM core proteins was retrieved (i) from ref.^9^, manually updated (DisGeNET, www.disgenet.com; OMIM disease terms MIM-phenotypes, www.omim.org) and categorized; (ii) from UniProt; (iii) from OpenTarget (platform.opentargets.org, filtered for ‘Genomics England PanelApp’ as source) and (iv) from the ‘Genomics England PanelApp’ (panelapp.genomicsengland.co.uk) as an additional source and quality filter (Supplementary Table 3). Only disease-links validated by Genomics England PanelApp (levels green and amber) were considered for further statistical evaluation.

### Protein quantification and profile building

For label-free quantification of proteins, LC-MS data were processed as described^28,88^. Briefly, m/z-calibrated precursor ion mass traces were extracted from MS raw files using MaxQuant v2.4.1 (www.maxquant.org) and mapped to the identified peptides using in-house software, including retention-time alignment and match-between-runs enabled (m/z window ±3 ppm, elution time window ±2 min). To account for technical variances between runs, intensities were corrected by first selecting for each run/slice up to 1024 peptides that have the lowest CV among all peptides with no missing values in a window of up to 8 slices before and after the current slice, then dividing the intensity values in that window by their mean and finally using the median of all relative intensities overlapping with the central slice as correction factor. Protein quantification and profile building were then carried out as previously described^28^, except that for protein quantification, peptide intensities from 7-11 consecutive slices were used (depending on the number of available peptide intensities). In a few cases of small proteins (e.g. hydrophobic MIC10 with two peptides), the protein abundance can be underestimated.

### Apparent molecular mass scaling

For mapping the migration distance on the blue native gel (i.e. slice index) to an apparent molecular mass, the theoretical protein masses of selected marker proteins/complexes were calculated using the respective full canonical sequence(s) in UniProt, and their log10 mass values were plotted versus the slice index of the respective profile-peak maxima. Apparent mass scales were obtained by least-squares-fitting an exponential function to the data (Extended Data Fig. 1b). Marker proteins/complexes were selected that (i) exhibit well-defined mono-disperse peaks in their profile(s), (ii) cover the entire range of slices analyzed, and (iii) exhibit well-established subunit composition/stoichiometry (as annotated in the PDB database or the UniProt curated information).

### Complexome data evaluation, profile peak complexity

The complexity estimation of individual protein abundance-mass profiles, i.e. the number of distinguishable peaks and shoulders that reflect one or multiple molecular entities, was based on a custom peak detection algorithm followed by fitting of a Gaussian mixture model. Potential peak locations, even inside peak shoulders, were identified by analyzing second derivatives and used to guide and constrain the model fitting. Accordingly, protein profiles were assigned ‘low’ complexity (1–4 components), ‘intermediate’ complexity (5–9 components), ‘high’ complexity (10–14 components) and ‘very high’ complexity (>15 components).

### Prediction of pairwise and preferred assembly of proteins

A convolutional neural network binary classifier (CNN) originally trained in-house with a large set of protein abundance over mass profile pairs from a mouse brain complexome dataset (3,300 pairs manually labelled positive, 9,900 pairs manually labelled negative, augmented and supplemented with random-sampled pairs to a training set of 609,120 pairs (positive:negative class ratio of 1:12)) was adapted to the HEK293T complexome dataset by transfer learning. The input for each candidate pair consisted of the two respective protein abundance profiles, the slice index-assigned molecular-weight scale, along with corresponding protein features such as protein molecular weights and the number of identified protein-specific peptides. Before training and inference, protein abundance profiles were processed to reduce noise and local profile gaps, and then resampled to a fixed representation of 120 data points. A set of manually labelled HEK complexome protein pairs (1950 positive, 6987 negative) was established for re-training. From these, training and validation subsets were generated with balanced representation of pairs (to avoid domination of individual multiprotein assemblies). Additional negative pairs were sampled from unlabeled candidate pairs under predefined constrains (protein naming, profile-overlap, abundance ratio). The CNN classifier was then re-trained by keeping most parameters fixed while unfreezing the second and third convolutional blocks together with the classification layers. Model training was performed with early stopping based on validation loss (binary cross entropy), and the final selected model corresponded to the checkpoint with the best validation performance on the HEK subset. The selected transfer-learned model was then applied to score all pairwise combinations of hMITCOM proteins, and finally evaluated based on the manually labelled subset (1111 positive, 1015 negative) as reference (Extended Data Fig. 2). Predicted interaction scores (independent of underlying interactions being direct or indirect) were used to rank candidate protein-protein relations for exploratory analysis in the complexome viewer or visualization of preferred protein assembly groups by t-SNE (Extended Data Fig. 3 and insets in the main figures showing preferred assembly groups of proteins).

### Tris-Tricine gel electrophoresis

For protein separation using 10% Tris-Tricine-SDS-PAGE, protein samples were mixed with Laemmli sample buffer with DTT and denatured at 65°C for 15 min. Electrophoresis was performed using cathode buffer (0.1 M Tris, 0.1 M Tricine, 0.1% [w/v] SDS, pH 8.25) and anode buffer (0.2 M Tris-HCl, pH 8.9) at 70 mA constant current. Subsequently, proteins were transferred to PVDF membranes (Immobilon-P; Millipore) in transfer buffer (20 mM Tris, 150 mM glycine, 0.02% [w/v] SDS, 20% [v/v]) ethanol) at 250 mA using the semi-dry system. Membranes were stained (40% [v/v] ethanol, 7% [v/v] acetic acid, 0.2% [w/v] Coomassie R250) and destained (30% [v/v] ethanol, 10% [v/v] acetic acid) until protein bands became visible. Membranes were subsequently cut, destained completely with methanol and blocked in blotting buffer (5% [w/v] skim milk powder, 20 mM Tris-HCl, pH 7.5, 125 mM NaCl) for 1 hr. The primary antibodies were incubated for 1 hr at room temperature or overnight at 4°C. Membranes were washed, incubated for 45 min with a peroxidase-coupled secondary antibody and developed with ECL developing solution using LAS-3000/4000 camera systems (Fujifilm/GE healthcare). The antibodies used are listed in Supplementary Table 5.

### TwinStrep affinity purification of protein complexes

TwinStrep affinity purification was carried out using the StrepTactin®XT system (IBA-Lifesciences). Either stably transfected cells or isolated mitochondria were solubilized in solubilization buffer (1% digitonin [w/v], 100 mM Tris-HCl pH 8, 150 mM NaCl, 10% glycerol, 1 mM PMSF, 1x protease inhibitor cocktail, 1000 U DNase I (Roche) for 30 min at 4°C. The solubilized cells or mitochondria were centrifuged at 18,000 x g to remove insolubilized material and the supernatant was moved to a fresh tube. StrepTactin®XT®4Flow (IBA-Lifesciences) resin was added to the supernatant according to the manufacturer protocol and incubated head-over-head, for 2 hrs or overnight at 4°C. Subsequently, the beads were moved to a polypropylene column and washed with 8 column volumes of washing buffer (0.3% digitonin [w/v], 100 mM Tris-HCl pH 8, 150 mM NaCl, 10% [v/v] glycerol, 1 mM PMSF). Elution was performed with 250 µl elution buffer (0.3% [w/v] digitonin, 100 mM Tris-HCl pH 8, 150 mM NaCl, 1 mM EDTA, 50 mM Biotin, 10% [v/v] glycerol, 1 mM PMSF) for 30 min at 4°C. The elution was either subjected to mass spectrometry or SDS-/blue native-PAGE.

### Affinity purification-mass spectrometry (AP-MS) analysis

Eluates from affinity-purifications were processed as described^89^. Dried peptide digests were dissolved in 13 µl 0.5% [v/v] trifluoroacetic acid each. 0.2 µl aliquots were loaded onto a C18 PepMap100 precolumn (300 µm i.d. × 5 mm; particle size 5 µm) with 0.05% [v/v] trifluoroacetic acid (5 min 20 µl/min) via the autosampler of a split-free UltiMate 3000 RSLCnano HPLC system (Thermo Fisher Scientific). Peptides were eluted using an aqueous-organic gradient (eluent A: 0.5% [v/v] acetic acid; eluent B: 0.5% [v/v] acetic acid in 80% [v/v] acetonitrile): 5 min 3% B, 60 min from 3% to 30% B, 10 min from 30% to 40% B, 5 min from 40% to 50% B, 5 min from 50% to 99% B, 5 min 99% B, 5 min from 99% to 3% B, 10 min 3% B (flow rate of 300 nl/min). Peptides were separated on a self pack nanoLC column (i.d. 75 µm; tip 8 µm; CoAnn Technologies) manually packed (25 cm) with ReproSil-Pur 120 ODS-3 (C18; particle size 3 µm; Dr Maisch HPLC) directly electrospraying at 2.3 kV (transfer capillary temperature 300°C) in positive ion mode into a Q Exactive HF-X mass spectrometer with a Nanospray Flex ion source (Thermo Fisher Scientific). Full MS instrument settings were: resolution 240,000, AGC target 3,000,000, maximum injection time 100 ms, scan range 370 to 1,700 m/z. Data-dependent MS/MS instrument settings were: resolution 15,000, AGC target 100,000, maximum injection time 200 ms, loop count 10 (TopN 10), isolation window 1.0 m/z, fixed first mass 100.0 m/z, normalized collision energy 28, minimum AGC target 8,000 (intensity threshold 40,000), charge exclusion unassigned, 1, and >5, peptide match preferred, exclude isotopes on, dynamic exclusion 30.0 s. m/z 445.12003 (protonated (Si(CH_3_)_2_O))_6_ from ambient air) was used as a lock mass^90^. Quantification of proteins was done as described above. For identification of C15orf61-associated proteins in APs versus controls, log10 values of the respective protein abundance ratios were normalized with respect to enrichment of the target (=1) and the mean distribution of background protein ratios (=0) yielding target-normalized ratios^91^ (tnRs) as a normalized measure of target-specific co-purification. Likewise, target-normalized abundance ratios (tnA values) were determined as a measure of co-purification efficiency (relative to C15orf61). The tnR/tnA-data were inspected by t-SNE (Extended Data Fig. 5a) generated in BELKI (https://github.com/phys2/belki; perplexity set to 50).

### Carbonate extraction

250 µg of purified mitochondria (protein amount) were incubated with 200 µl of 100 mM sodium carbonate at either pH 11.5 or pH 10.5 for 30 min on ice. All samples were subsequently centrifuged for 1 hr at 100,000 x g at 4°C. The supernatants were collected and proteins were precipitated using 72% TCA (20% [v/v]) and 2% sodium desoxycholate (1% [v/v]). The pellets were washed with ice-cold unbuffered Tris (1M), incubated with 2x Laemmli sample buffer at 60°C for 10 min and analyzed by SDS-PAGE followed by immunodecoration.

### Preparation for FLAG immunoisolation

Human HEK293-Flp-In cells (Thermo Fisher) were cultured in high glucose (4,5 mg/ml) containing DMEM (Dulbecco’s modified Eagle’s medium), supplemented with 10% (v/v) FBS (fetal bovine serum) (Capricorn Scientific), 1 mM sodium pyruvate, 2 mM L-glutamine under a humidified 5% CO2 containing atmosphere at 37°C.

TIM21^FLAG^ (ref.^79^) or ^FLAG^OXA1L (ref.^74^) were expressed in HEK293 cells by addition of tetracycline (final 0,1 µg/ml) for 24h. Cells were harvested with PBS and mitochondria isolated with TH-buffer^43^. In brief, cells were harvested after 24h induction using PBS. After pelleting (1250 x g, 5 min, 4°C), mitochondria were extracted by homogenisation in TH-buffer (300 mM trehalose, 10 mM KCl, 10 mM HEPES, 0.1 mg/ml BSA (Sigma-Aldrich, A6003-100G), pH 7.4) using a Dounce Homogenizer. To remove non-homogenized cells, the material was centrifuged at 400 xg, 4°C for 10 min. These steps were repeated once, followed by an additional clearing spin at 800 xg, 4°C, 5 min. Afterwards, mitochondria were pelleted at 10,000 xg, 4°C for 10 min and washed once with BSA-free TH buffer. Prior to the Bradford assay, mitochondria were pelleted again and resuspended in BSA-free TH buffer.

For FLAG-immunoisolation, TIM21^FLAG^ or ^FLAG^OXA1L mitochondria were suspended in isolation buffer (50 mM Tris/HCl pH 7.4; 150 mM NaCl; 20 mM MgCl_2_; 10% glycerol; 1 mM PMSF and 1% digitonin) and kept under mild agitation for 30 min at 4°C. Debris was removed by centrifugation (16,000 x g at 4°C, 15 min) and supernatants transferred to anti-FLAG M2 Affinity Gel (Sigma), incubated by rotating at 4°C for 1h, prior to washing with wash buffer (50 mM Tris/HCl pH 7.4; 150 mM NaCl; 20 mM MgCl_2_; 10% glycerol; 1 mM PMSF and 0.3% digitonin). Proteins were eluted by competition with FLAG peptides and applied to western blot analysis.

### Preparation for standard blue native analysis

HEK293T cells were harvested, washed in PBS, and resuspended in ice-cold Buffer A (83 mM Sucrose, 10 mM HEPES, pH 7.2). After homogenizing the cells with 20 strokes in a Dounce homogenizer, an equal volume of buffer B (250 mM Sucrose, 30 mM HEPES, pH 7.2) was added, and the homogenate was centrifuged for 5 min at 1,000 x g at 4°C. The resulting supernatant was centrifuged again at 12,000 x g for 10 min at 4°C and the mitochondrial pellet was resuspended in Buffer C (320 mM Sucrose, 1 mM EDTA, 10 mM Tris-Cl pH 7.4). Protein concentration was determined using Roti®-Quant.

Isolated mitochondria were pelleted at 15,000 x g for 5 min at 4°C and gently resuspended in digitonin buffer (1% [w/v] digitonin, 20 mM Tris-HCl pH 7.4, 0.1 mM EDTA, 50 mM NaCl, 10% [v/v] glycerol, and 1 mM PMSF). After solubilization for 30 min on ice, non-solubilized material was removed by a clarifying spin at 15,000 x g for 10 min at 4°C. The supernatant was mixed with loading dye (5 % Coomassie blue G, 500 mM ε-amino n-caproic acid in 100 mM Bis Tris pH 7.0) and samples directly loaded onto either a 4-13% or 5-10% blue native-PAGE, depending the resolution required for the protein complexes. Protein complexes were analyzed by immunodecoration.

### Protein complex modeling

The structure of the SAM50a/b-MTX2-MTX3 protein complex was predicted using AlphaFold3 (ref.^92^). For each protein, the canonical isoform sequence was submitted. All five generated models converged on highly similar conformations for both complexes, indicating robust and consistent predictions. The top-ranked models were selected for further analysis and exported to PyMOL Molecular Graphics System (Version 3.0, Schrödinger LLC) for figure preparation. The partially disordered region of MTX3 (Δ284–312) exhibiting poor local confidence scores (pLDDT <70) was excluded from the structural representation.

## Acknowledgements

We thank Florian Wollweber for technical support and discussion. Work included in this study has also been performed in partial fulfilment of the requirements for the doctoral thesis of A.F. (University of Freiburg). This work was supported by the Deutsche Forschungsgemeinschaft (DFG, German Research Foundation) PF 202/9-1 project ID 394024777 (to N.P.), SFB 1381 project ID 403222702 (to U.S., B.F. and N.W.), FOR 2848 project ID 401510699 (to M.v.d.L. and H.R.), SPP 2453 project IDs 541555098 (to T.B.) 541758477 (to M.v.d.L.) 541758684 (to S.D.), SFB 1218 project ID 269925409, BE 4679/9-1 project ID 528247081, BE 4679/13-1 project ID 568735694 (to T.B.), and Germany’s Excellence Strategy (CIBSS – EXC-2189 – project ID 390939984 to U.S., B.F., N.W. and N.P.).

## Author contributions

U.S., B.F., N.W. and N.P. conceived and supervised the project. U.S., A.H., C.S., W.B., J.B.v.d.B., C.S.M., H.R., B.F., N.W. and N.P. acquired and evaluated data related to BN-MS analyses. U.S., A.H. and B.F. set up hMitCOM for online access. C.S., L.M., A.F., K.v.d.M. and S.D. performed biochemical experiments, characterized complexes and analyzed the results together with U.S., T.B., M.v.d.L., H.R., B.F., N.W. and N.P.. U.S., A.H., C.S., L.M., A.F., K.v.d.M., S.D., H.R., B.F., N.W. and N.P. prepared figures. N.P., U.S., N.W. and B.F. wrote the manuscript with the support of all authors.

## Competing interests

The authors declare no competing interests.

**Correspondence and requests for materials** should be addressed to Nikolaus Pfanner, Nils Wiedemann or Bernd Fakler.

## Extended Data Figure Legends

**Extended Data Fig. 1.**
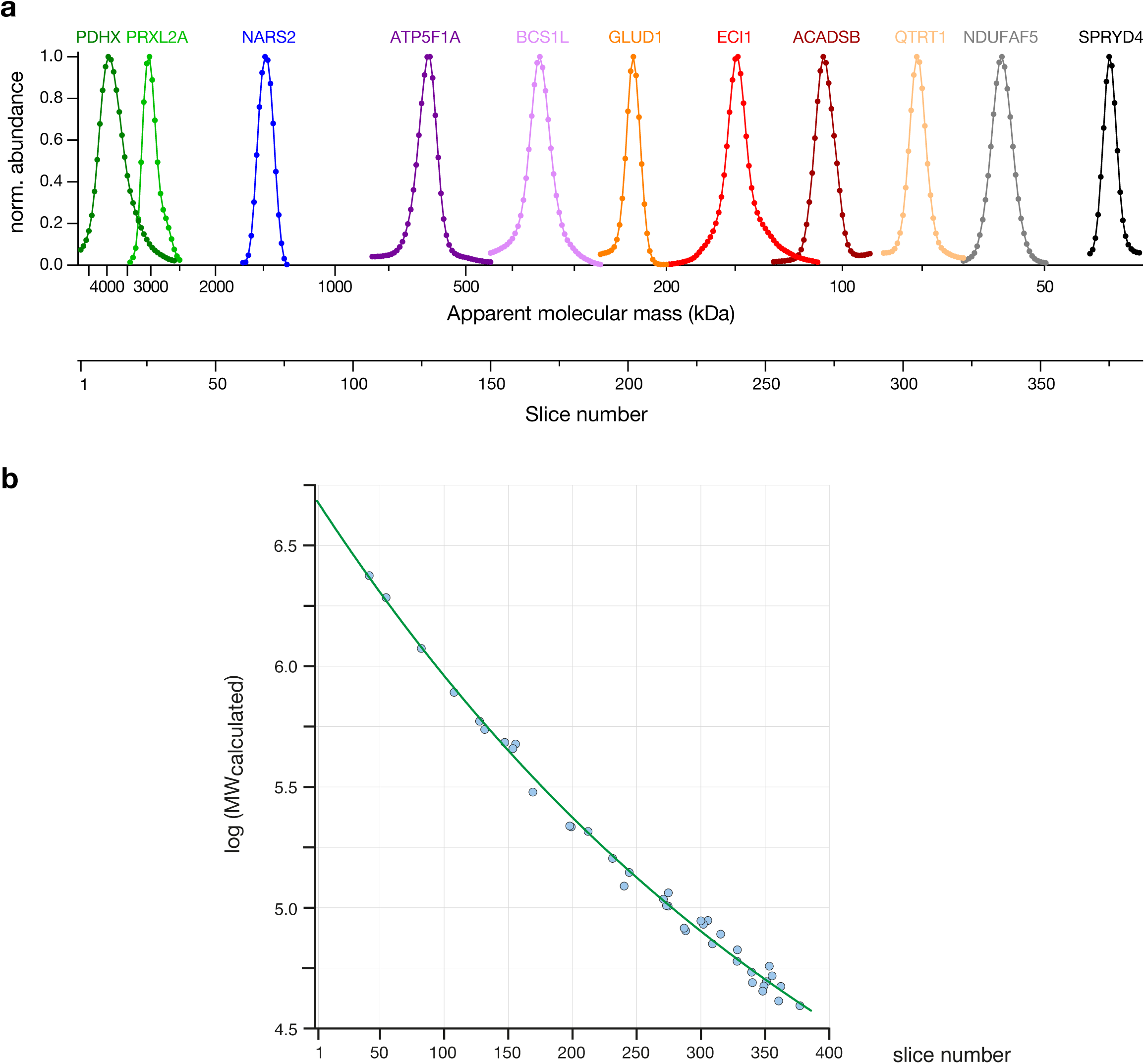
Benchmarking of hMitCOM profiles. **a**, Profile peaks of the indicated mitochondrial proteins, showing the resolution over the molecular mass range of the blue native gel. **b**, Conversion of slice index to apparent molecular mass. Plot shows selected marker proteins or protein complexes (circles in blue) with derived calculated Log(predicted molecular mass, MW): PDD2L, SMYD5, BCAT2, PDP2, GATA, DBLOH dimer, PPA6, TMLH, QCR1, SYEM, PCKGM, FOLC, ACSL1, CPT2, QTRT2+TGT heterodimer, SUV3, AADAT dimer, NEUL, ACON, AL1L2, SYVM, C1TM, SYIM, SYVC, SDHA/II, ABCB7 homodimer, C1TM homodimer, COX1/IV, NNTM homodimer, IDH3A heterooctamer, QCR1/III_2_, UGPA homooctamer, ACLY homotetramer, FAS homodimer, ATPA/V, PCCB heterohexamer, ATPA/V_2_, COX1/I_1_III_2_IV_2_, ATPG/V_4_. Green line is a mono-exponential fit to the data used for converting slice number scale into apparent molecular mass scale.

**Extended Data Fig. 2.**
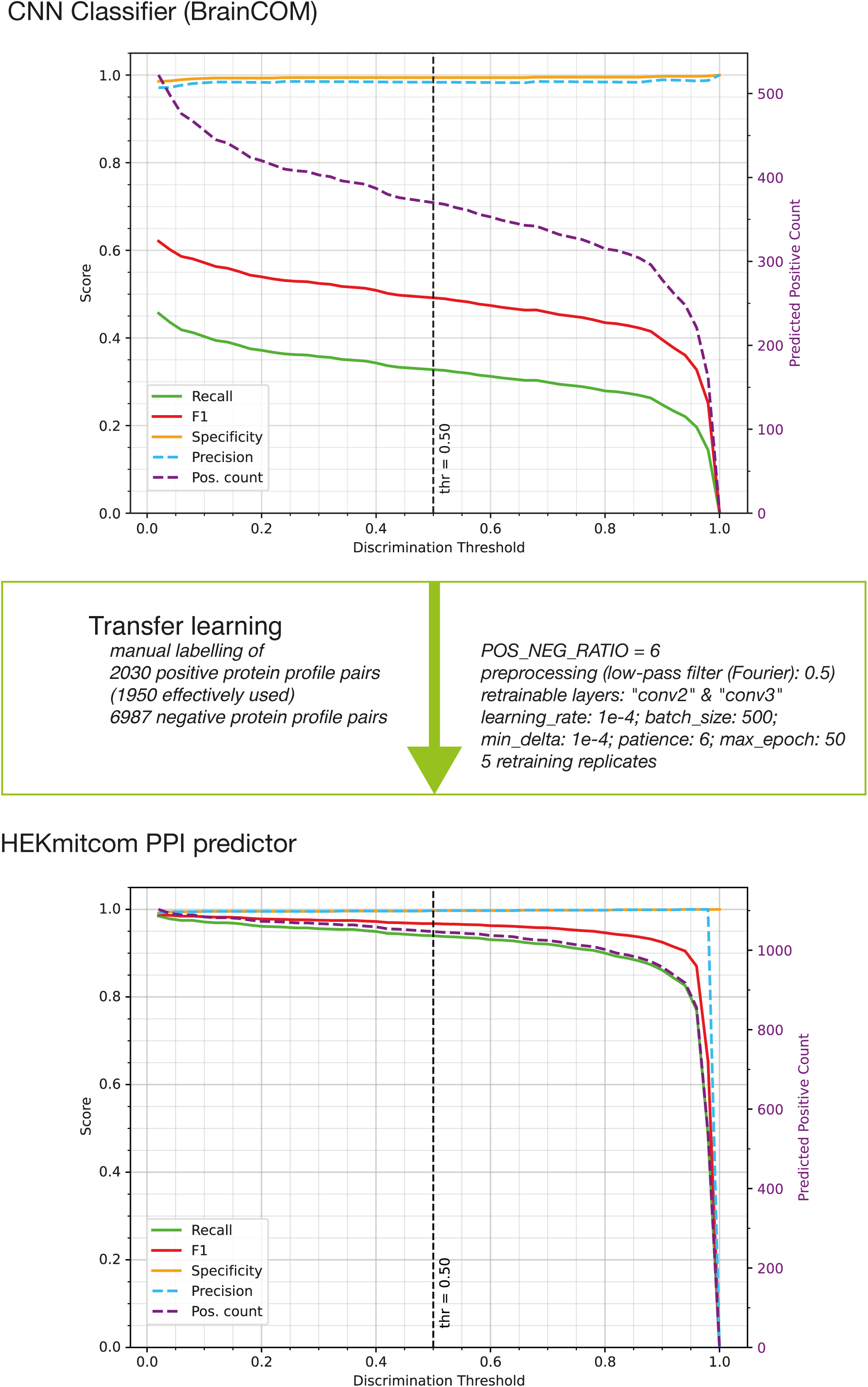
Performance of the CNN classifier (PPI Finder) upon transfer learning. Plot showing the performance of PPI-Finder (a CNN trained classifier to identify protein profile pairs indicating protein-protein interactions; Methods) before (**a**) and after (**b**) transfer learning to optimally adapt to the hMitCOM dataset using manually labelled positive (n=1111) and negative (n=1052) hMitCOM protein profile pairs as reference. Validation metrics (at default prediction score threshold of 0.5) in (a) were positive recall (sensitivity): 32.8% (370 predicted labelled positive pairs), precision: 98.4%, specificity: 99.4%, F1 score (weighted harmonic mean of precision and recall): 49.2%. Note that model re-training strongly enhanced the performance of the classifier finally used for hMitCOM (b): positive recall (sensitivity) was 94% (1047 predicted labelled positive pairs), precision was 99.7%, specificity was 99.7%. The F1 score (96.8%) was used as target for transfer learning optimization.

**Extended Data Fig. 3.**
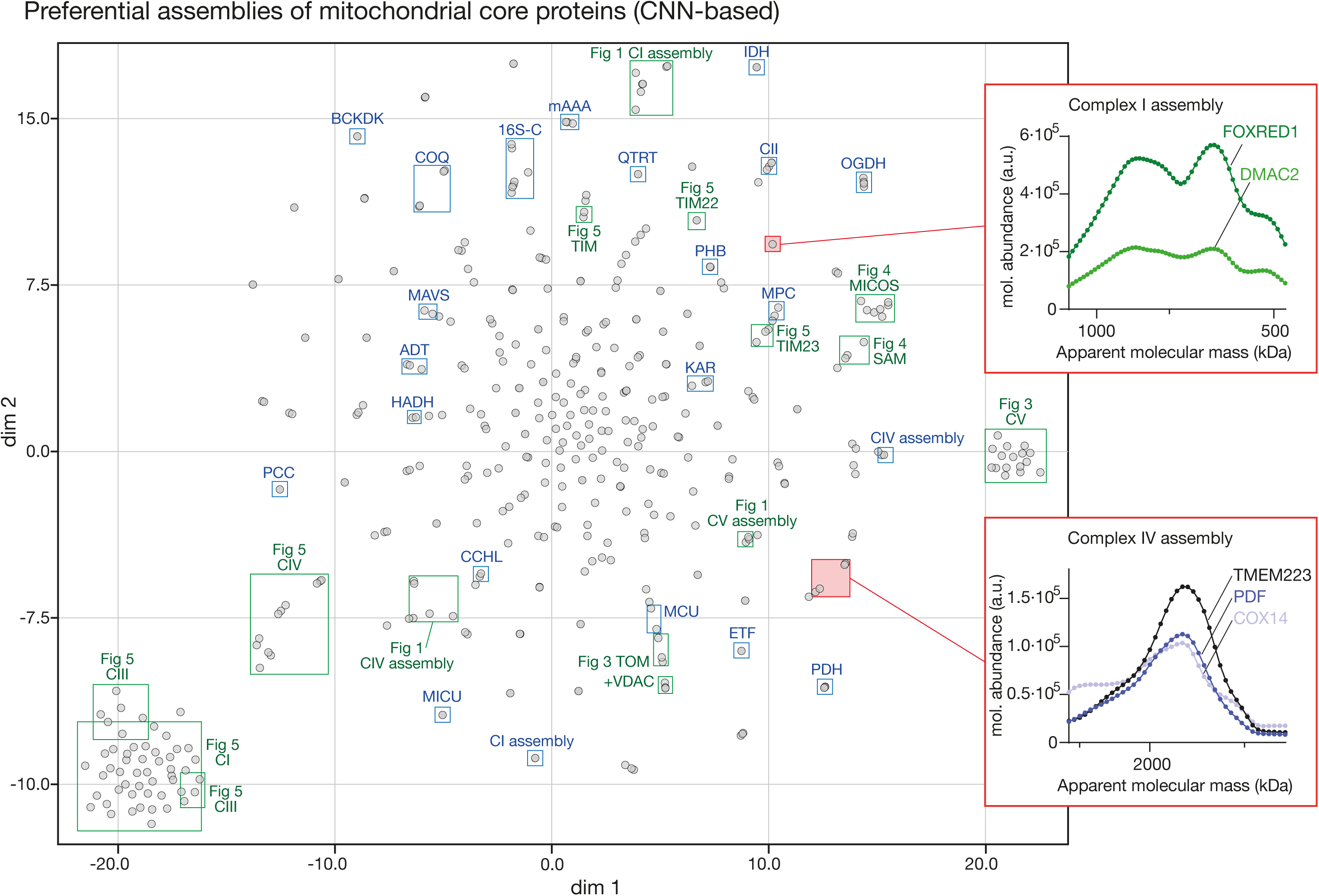
Preferential assemblies of hMitCOM proteins based on AI-derived pairwise PPI-scores. t-SNE plot showing similarity-based clustering of hMitCOM core proteins (392 with PPI scores >0.01) into preferred assemblies comprising one or more protein complexes. Blue boxes highlight assemblies/complexes annotated in UniProt, green boxes mark protein assemblies described by their profiles in the indicated Figures. Two examples of novel assemblies formed by Complex I and Complex IV assembly factors, respectively, are framed red and shown in insets as zooms into congruent regions of their abundance-mass profiles.

**Extended Data Fig. 4.**
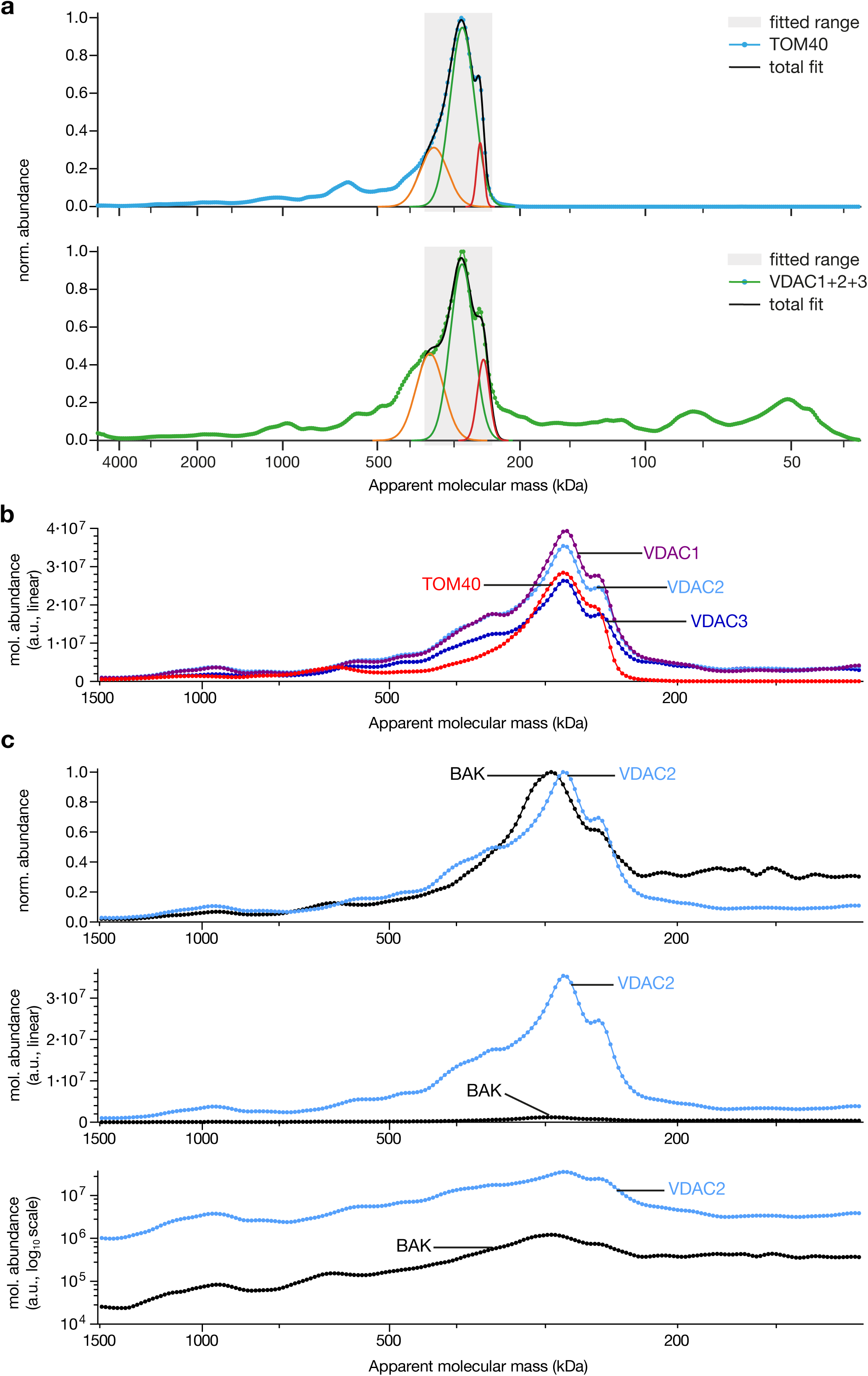
Comparison of TOM40, VDAC and BAK profiles. **a**, Quantification of TOM40 co-migration with VDACs. Multi-gaussian fits (black lines) to a central region (grey) of abundance-mass profiles of TOM40 (upper panel, blue) and total VDACs (i.e. sum of VDAC1, VDAC2, VDAC3 abundance-mass profiles; lower panel, green) revealed at least three common peak components (orange, red and green gaussians). Co-migration in joint assemblies (total area under profile minus area under the fitted components) can be estimated to be at least 75% for TOM40 and 45% for VDACs. **b**, Molecular (mol.) abundance-mass profiles of TOM40, VDAC1, VDAC2 and VDAC3. **c**, Comparison of normalized and molecular abundance-mass profiles of BAK and VDAC2.

**Extended Data Fig. 5.**
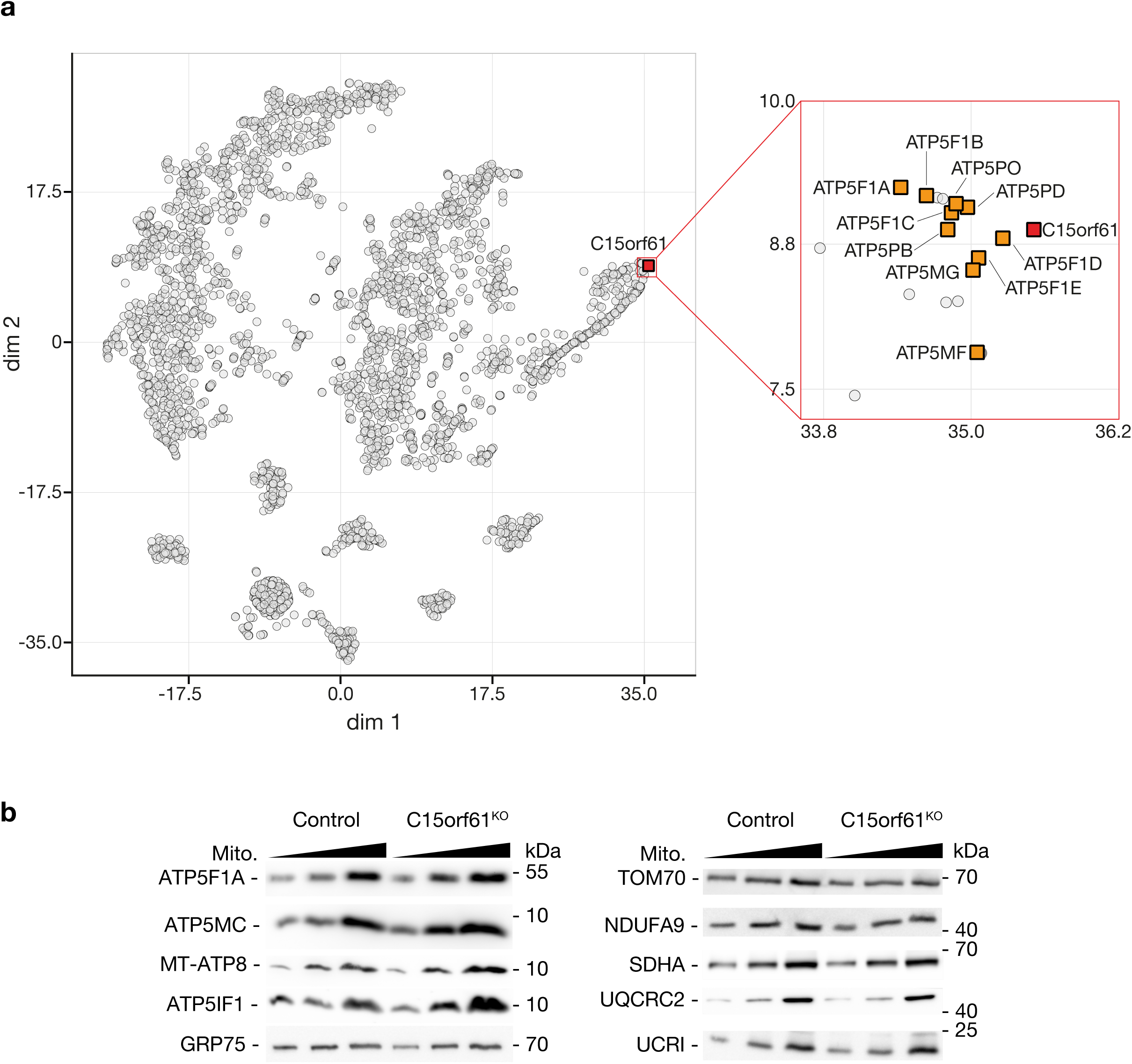
Characterization of C15orf61. **a**, Affinity purification-mass spectrometry (AP-MS) analysis of tagged C15orf61. t-SNE plot (proteins represented by circles) showing the result of three replicate C15orf61 AP-MS analyses, based on target-normalized ratios (tnR values) and target-normalized abundances (tnA values). Inset shows the vicinity of C15orf61 at enlarged scale with subunits of F_1_F_o_-ATP synthase found as the closest (i.e. most specifically and abundantly co-purified) interaction partners of C15orf61. **b**, Mitochondria (Mito.) isolated from either wild-type (WT) or C15orf61^KO^ cells were subjected to SDS-PAGE followed by immunodecoration with the indicated antibodies.

**Extended Data Fig. 6.**
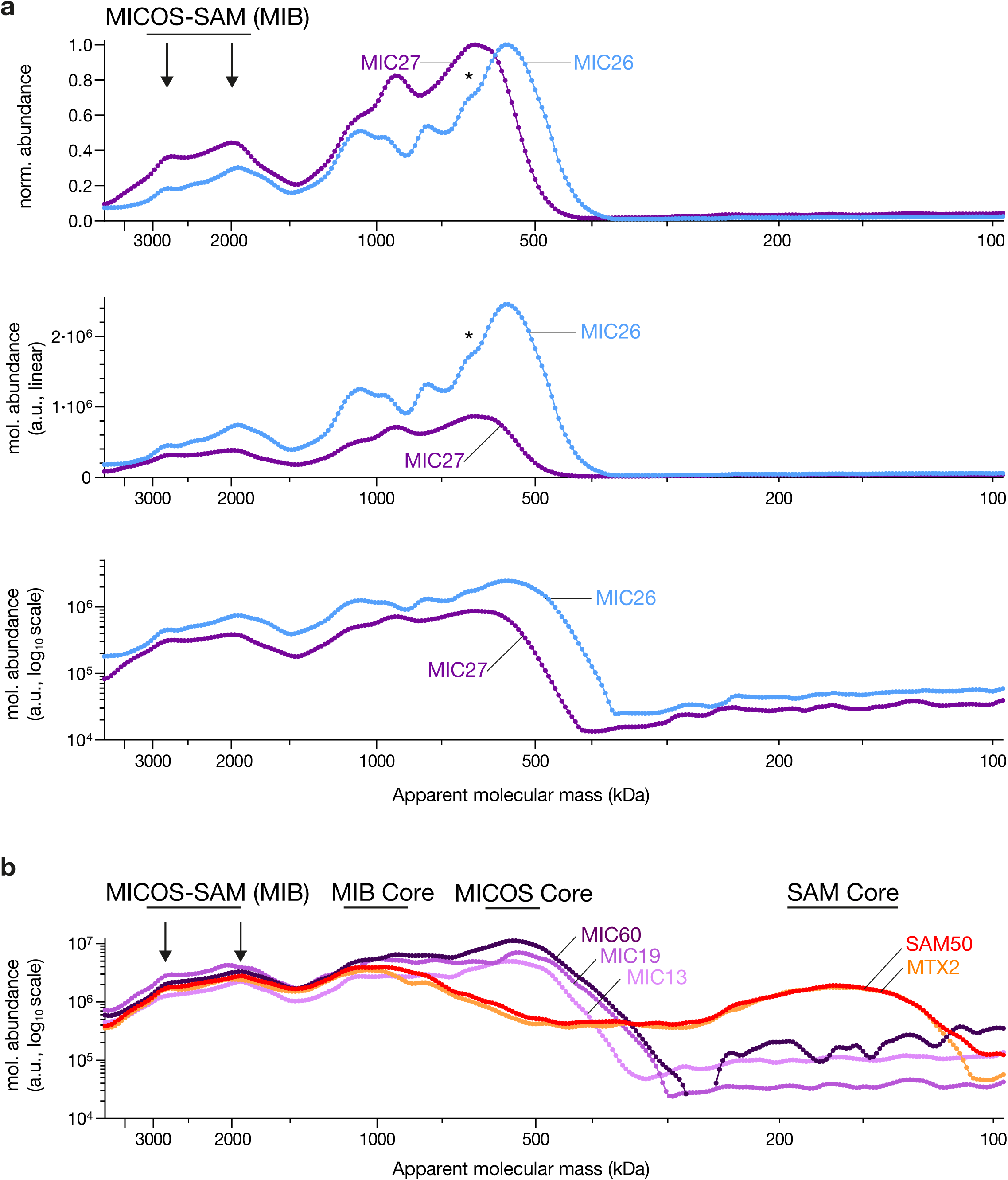
Characterization of MICOS, SAM and supercomplexes. **a**, Comparison of normalized (norm.) and molecular (mol.) abundance-mass profiles of MIC26 and MIC27. Asterisk, shoulder of the MIC26 profile aligning with the major MIC27 peak. **b**, Molecular abundance-mass profiles of MICOS and SAM subunits. MICOS-SAM (MIB) supercomplexes, MIB Core, MICOS Core and SAM Core are indicated.

**Extended Data Fig. 7.**
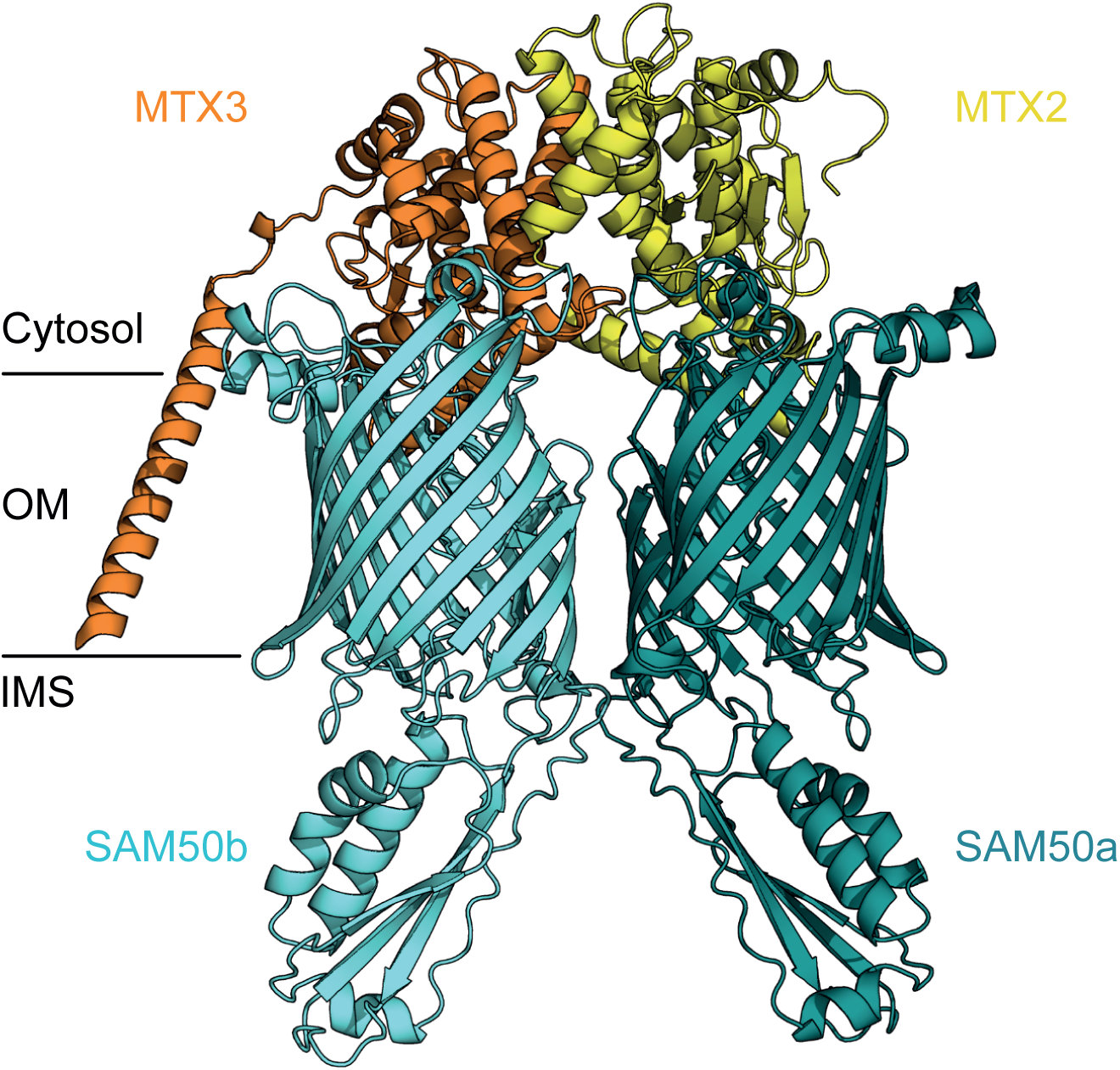
Model of human SAM complex. AlphaFold3 protein complex model of a human SAM heterotetramer consisting of MTX3 (orange), MTX2 (yellow), SAM50b (cyan) and SAM50a (dark teal) (ipTM = 0.52, pTM = 0.58), resembling the yeast SAM complex (Sam37/35/50b and 50a)^93^. Predicted structure in cartoon representation is viewed parallel to the membrane plane. The lines indicate the predicted membrane region (OM, outer mitochondrial membrane); IMS, intermembrane space. According to yeast data, SAM50-MTX2-MTX3 represents the functional SAM complex (Takeda-2021), however, in human mitochondria these proteins are not observed together in the SAM core complex of ∼160 kDa, but only in MICOS-SAM (MIB) supercomplexes of 1 MDa and larger (see also Fig. 4e).

## Supplementary Information Guide

Uwe Schulte et al.

**Supplementary Table 1. Data underlying abundance-mass profiles of hMitCOM proteins (<u>separate Excel file</u>).**

**Supplementary Table 2. Annotation of hMitCOM proteins (<u>separate Excel file</u>).**

**Supplementary Table 3. Disease association of hMitCOM genes (<u>separate Excel file</u>).**

---

**Supplementary Table 4.**
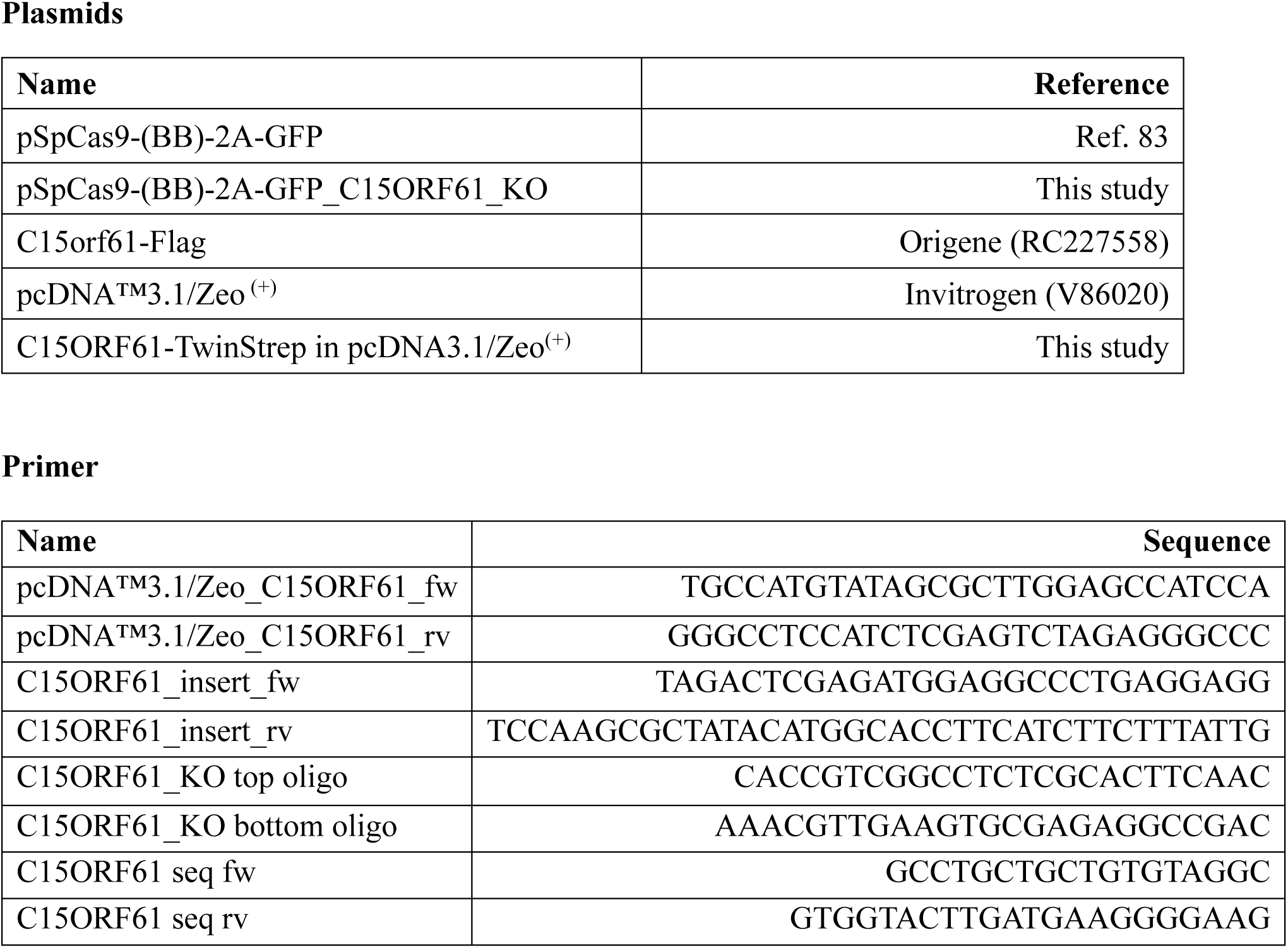
Plasmids and primers used in this study.

**Supplementary Table 5.**
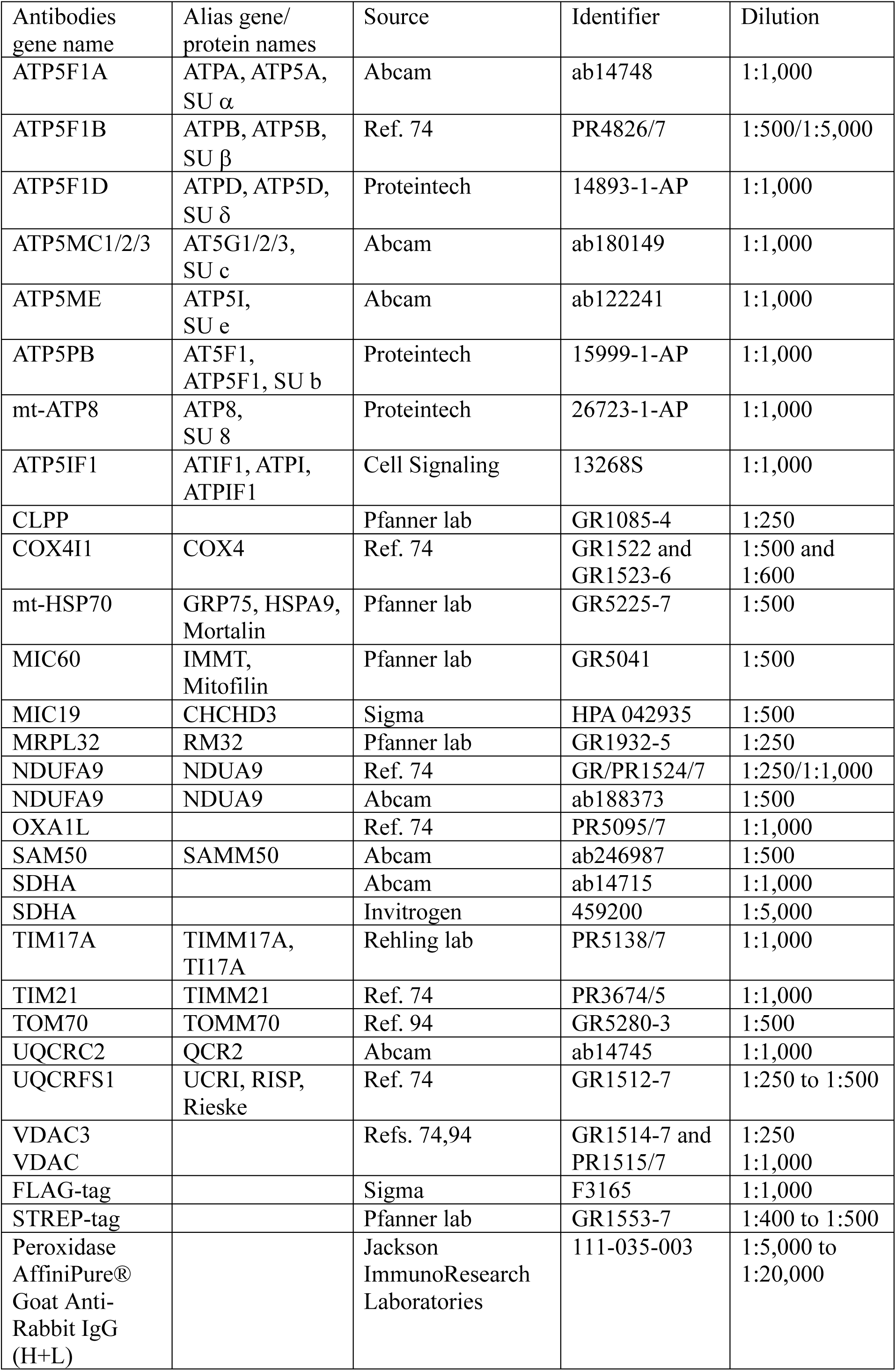
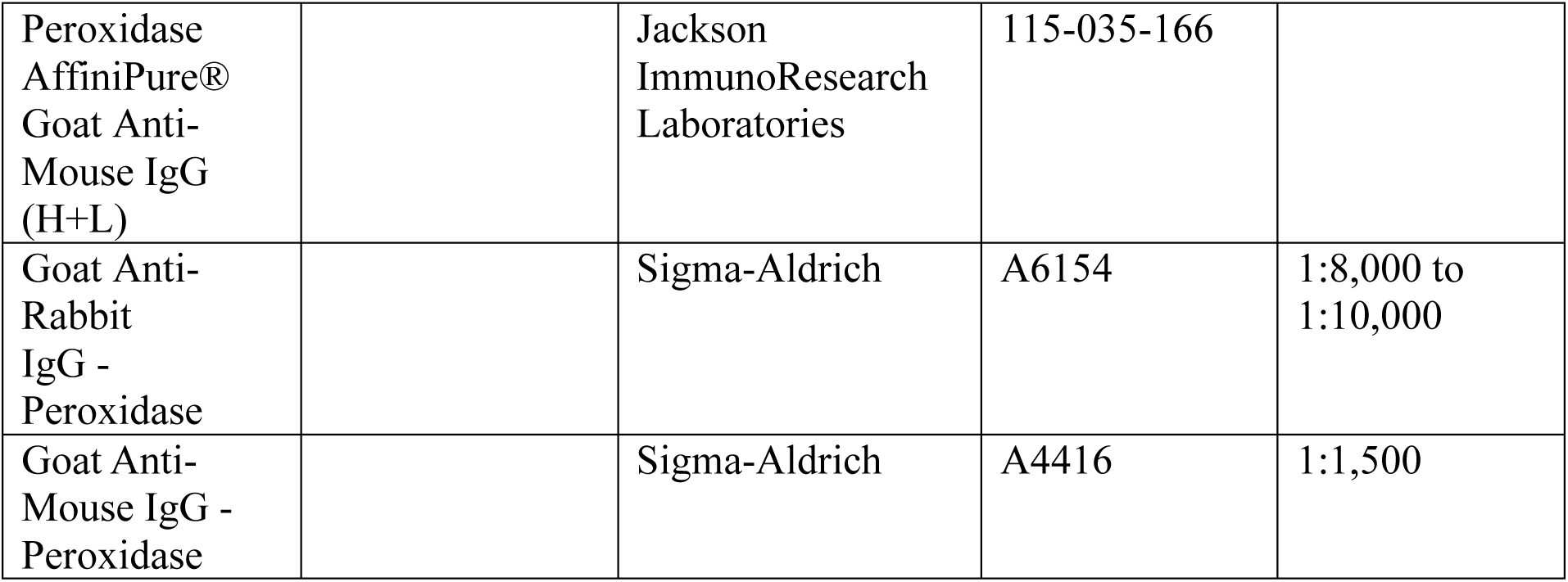
Antibodies used in this study.

